# A SARS-CoV-2 neutralizing antibody selected from COVID-19 patients by phage display is binding to the ACE2-RBD interface and is tolerant to most known recently emerging RBD mutations

**DOI:** 10.1101/2020.12.03.409318

**Authors:** Federico Bertoglio, Viola Fühner, Maximilian Ruschig, Philip Alexander Heine, Leila Abasi, Thomas Klünemann, Ulfert Rand, Doris Meier, Nora Langreder, Stephan Steinke, Rico Ballmann, Kai-Thomas Schneider, Kristian Daniel Ralph Roth, Philipp Kuhn, Peggy Riese, Dorina Schäckermann, Janin Korn, Allan Koch, M. Zeeshan Chaudhry, Kathrin Eschke, Yeonsu Kim, Susanne Zock-Emmenthal, Marlies Becker, Margitta Scholz, Gustavo Marçal Schmidt Garcia Moreira, Esther Veronika Wenzel, Giulio Russo, Hendrikus S.P. Garritsen, Sebastian Casu, Andreas Gerstner, Günter Roth, Julia Adler, Jakob Trimpert, Andreas Hermann, Thomas Schirrmann, Stefan Dübel, André Frenzel, Joop Van den Heuvel, Luka Čičin-Šain, Maren Schubert, Michael Hust

**Author notes:** shared first authors (Federico Bertoglio, Viola Fühner, Maximilian Ruschig, Philip Alexander Heine, Leila Abasi, Thomas Klünemann, Ulfert Rand). shared last authors (André Frenzel, Joop Van den Heuvel, Luka Čičin-Šain, Maren Schubert, Michael Hust). Corresponding author for antibody development: Michael Hust Technische Universität Braunschweig Institut für Biochemie, Biotechnologie und Bioinformatik Abteilung Biotechnologie Spielmannstr. 7 38106 Braunschweig, Germany. Corresponding author for antigen production: Maren Schubert Technische Universität Braunschweig Institut für Biochemie, Biotechnologie und Bioinformatik Abteilung Biotechnologie Spielmannstr. 7 38106 Braunschweig, Germany. Corresponding author for virus neutralization: Luka Ĉicin-Ŝain Helmholtz Centre for Infection Research (HZI) Department of Vaccinology and Applied Microbiology, Inhoffenstr. 7 38124 Braunschweig, Germany. Corresponding author for crystallography: Joop van den Heuvel Helmholtz Centre for Infection Research (HZI) Department of Structure and Function of Proteins Inhoffenstr. 7 38124 Braunschweig Germany. Corresponding author for preclinical development of STE90-C11 (COR-101): André Frenzel CORAT Therapeutics GmbH Science Campus Braunschweig-Süd Inhoffenstr. 7 38124 Braunschweig Germany.

## Abstract

The novel betacoranavirus SARS-CoV-2 causes a form of severe pneumonia disease, termed COVID-19 (coronavirus disease 2019). Recombinant human antibodies are proven potent neutralizers of viruses and can block the interaction of viral surface proteins with their host receptors. To develop neutralizing anti-SARS-CoV-2 antibodies, antibody gene libraries from convalescent COVID-19 patients were constructed and recombinant antibody fragments (scFv) against the receptor binding domain (RBD) of the S1 subunit of the viral spike (S) protein were selected by phage display. The selected antibodies were produced in the scFv-Fc format and 30 showed more than 80% inhibition of spike (S1-S2) binding to cells expressing ACE2, assessed by flow cytometry screening assay. The majority of these inhibiting antibodies are derived from the VH3-66 V-gene. The antibody STE90-C11 showed a sub nM IC50 in a plaque-based live SARS-CoV-2 neutralization assay. The *in vivo* efficacy of the antibody was demonstrated in the Syrian hamster and in the hACE2 mice model using a silenced human IgG1 Fc part. The crystal structure of STE90-C11 Fab in complex with SARS-CoV-2-RBD was solved at 2.0 Å resolution showing that the antibody binds at the same region as ACE2 to RBD. The binding and inhibtion of STE90-C11 is not blocked by many known RBD mutations including N439K, L452R, E484K or L452R+E484Q (emerging B.1.617). STE90-C11 derived human IgG1 with FcγR silenced Fc (COR-101) is currently undergoing Phase Ib/II clinical trials for the treatment of moderate to severe COVID-19.

**In Brief:** Human antibodies were selected from convalescent COVID-19 patients using antibody phage display. The antibody STE90-C11 is neutralizing authentic SARS-CoV-2 virus *in vitro* and *in vivo* and the crystal structure of STE90-C11 in complex with SARS-CoV-2-RBD revealed that this antibody is binding in the RBD-ACE2 interface. S1 binding of STE90-C11 and inhibition of ACE2 binding is not blocked by many known RBD mutations.

## Introduction

The severe pneumonia COVID-19 (coronavirus disease 2019) is a disease caused by the novel coronavirus SARS-CoV-2 and was described at the end of 2019 in Wuhan China (Lu et al., 2020; Zhou et al., 2020). This new human pathogenic coronavirus is closely related to the bat coronavirus RATG13 indicating an animal-to-human transition (Shang et al., 2020a). The spike (S) protein of SARS-CoV-2 binds to the human zinc peptidase angiotensin-converting enzyme 2 (ACE2) as main receptor for cell entry. The S protein has two subunits: the N-terminal S1 harbouring the receptor binding domain (RBD) and the viral membrane-anchored C-terminal S2 subunit, which is required for trimerization and fusion of the virus and host membrane for viral entry (Shang et al., 2020b; Starr et al., 2020; Walls et al., 2020; Wang et al., 2020b; Wrapp et al., 2020; Yan et al., 2020). Blocking of the RBD-ACE2 interaction by therapeutic antibodies as a strategy to treat COVID-19 (Zhou and Zhao, 2020) is a promising approach since the successful generation of neutralizing antibodies against S1 and RBD was already demonstrated for SARS-CoV and MERS (Coughlin and Prabhakar, 2012; Widjaja et al., 2019; Zhu et al., 2007). Some of the anti-SARS-CoV antibodies showed also cross-reaction and cross-neutralization of SARS-CoV-2 (Huo et al., 2020; Tian et al., 2020; Wang et al., 2020a).

Antibody phage display is a widely used *in vitro* technology to select human antibody fragments for the development of therapeutic antibodies (Frenzel et al., 2016). Currently, 14 EMA or FDA approved antibodies were generated by antibody phage display (Alfaleh et al., 2020). Human antibodies can be selected from two kind of antibody gene libraries: universal libraries and immune libraries. Universal libraries which have a naïve, semi-synthetic or synthetic antibody gene repertoire, allow – in theory - the selection of antibodies against any kind of molecule, while immune libraries derived from immunized donors typically facilitate the antibody generation against the pathogen that affected the donors (Bradbury et al., 2011; Frenzel et al., 2017; Kretzschmar and von Rüden, 2002; Kügler et al., 2018; Kuhn et al., 2016). Immune phage display libraries from convalescent patients or immunized human donors were used in the past for the successful generation of antibodies against infectious diseases (Duan et al., 2009; Rahumatullah et al., 2017; Trott et al., 2014; Wenzel et al., 2020a; Zhang et al., 2003). Human antibodies were also selected directly against SARS-CoV-2 using different approaches including single B-cell cloning or antibody phage display (Baum et al., 2020; Bertoglio et al., 2021; Kreer et al., 2020; Robbiani et al., 2020; Shi et al., 2020; Zeng et al., 2020).

Recently, several SARS-CoV-2 variants with mutations in the RBD emerged. Here most prominent are the B.1.1.7 (“UK”, RBD mutation N501Y), B.1.351 (“Southafrica”, K417N, E484K, N501Y) and P1 (B.1.1.28.1) (“Brazil”, K417T, E484K, N501Y) (Rees-Spear et al., 2021). More recently other variants like B.1.429+B.1.427 (“Southern California”, L452R) (Zhang et al., 2021), B.1.526 (“New York”, E484K, in some variants S477N instead of E484K) (Annavajhala et al., 2021), B.1.258Δ (“Czech”, N439K) (Brejová et al., 2021; Surleac et al., 2021), P2 (B.1.1.28.2) (E484K) (Nonaka et al., 2021), P3 (B.1.1.28.3) (E484K, N501Y) (Tablizo et al., 2021), B.1.1.33 (E484K) (Resende et al., 2021), B.1.617 (“India”, L452R, E484Q) (Ranjan et al., 2021) and other variants like B.1.525 (E484K) (Hodcroft et al., 2021) occurred.

In this work, immune phage display libraries from six COVID-19 convalescent patients were constructed, RBD-binding antibodies have been selected resulting in SARS-CoV-2 inhibiting and neutralizing antibodies. The crystal structure of the best neutralizing candidate STE90-C11 in complex with wt RBD was elucidated. Together with its unique binding pattern that tolerates a large number of RBD mutations, its properties suggest its use as a therapeutic agent against SARS-CoV-2 infections.

## Results

### Immune library construction

For the immune library construction, we collected blood samples of 16 q-RT-PCR confirmed COVID-19 convalescent patients. To analyze the immune answer, sera of all patients were titrated on RBD (Supplementary Data 1). The patients 1,2,5,6,9 and 14 showed the highest IgG titers against RBD and were chosen for library construction. PBMCs were isolated and CD19+/CD138+ plasma cells were enriched by FACS from some of these samples (Supplementary Data 2). Both PBMCs and plasma cells were used for library construction. Six individual libraries were constructed from PBMCs of patients 2, 6 and 14, while four additional libraries were made from CD19+/CD138+ plasma cells pools (pool 1: patients 2,6,14; pool 2: patients1,5,9). All libraries were constructed cloning kappa and lambda antibody genes separately, resulting in diversities ranging from 0.7×10 ^7^ to 1.7×10^8^ estimated independent clones.

### Antibody fragments were selected against SARS-CoV-2

Antibodies were selected by panning in microtiter plates against SARS-CoV-2 S1 subunit using S1-S2 in the first panning round, RBD in the second panning round and S1-S2 in the third panning round. This approach focused on the antibody selection on epitopes located within the RBD while being presented in the correct conformation of the intact spike protein trimer. The two plasma cell-derived immune libraries for kappa, respectively lambda, were combined and the panning was performed separately for lambda and kappa libraries. All kappa and lambda libraries originating from the PBMCs were pooled together. The following single clone screening was performed by antigen ELISA in 96 well MTPs, using soluble monoclonal scFv produced by *E. coli*. 542 monoclonal hits were identified. DNA sequencing revealed 197 unique antibodies. The V-Gene combinations for kappa (Supplementary Data 3) and lambda (Supplementary Data 4) antibodies were analyzed. The most abundantly used V-genes were VH3-66, VK1D-39 und VL3-21. Antibody sequences that showed potential developability liabilities, i.e. glycosylation sites or single cysteines in the CDRs were excluded from further analysis. 135 antibodies were re-cloned into the bivalent scFv-Fc format and produced in Expi293F cell in 5 mL culture scale, with yields ranging from of 70 to 260 mg/L.

### Screening for inhibition in a cell-based assay

To screen for SARS-CoV-2 blocking antibodies, an inhibition assay was performed by flow cytometry on ACE2-expressing EXPI293F cells, measuring direct competition between S1-S2 trimer and scFv-Fc antibodies. The entire spike protein ectodomain was used for this inhibition assay for optimal representation of the viral binding. In a first screening, the 135 antibodies were tested at 1500 nM S1-S2 (molar ratio antibody:S1-S2 30:1) (Figure 1). 30 antibodies that inhibited S1-S2 binding more than 80% were selected for further analysis. The germlines genes of these inhibiting scFv-Fc are given in Table 1. VH3-66 was the most abundantly used heavy chain V-gene of the well inhibiting scFv-Fc. All VH3-66 derived antibodies had closely related VH CDRs amino acid sequence while the corresponding light chains were different.

**Fig. 1.**
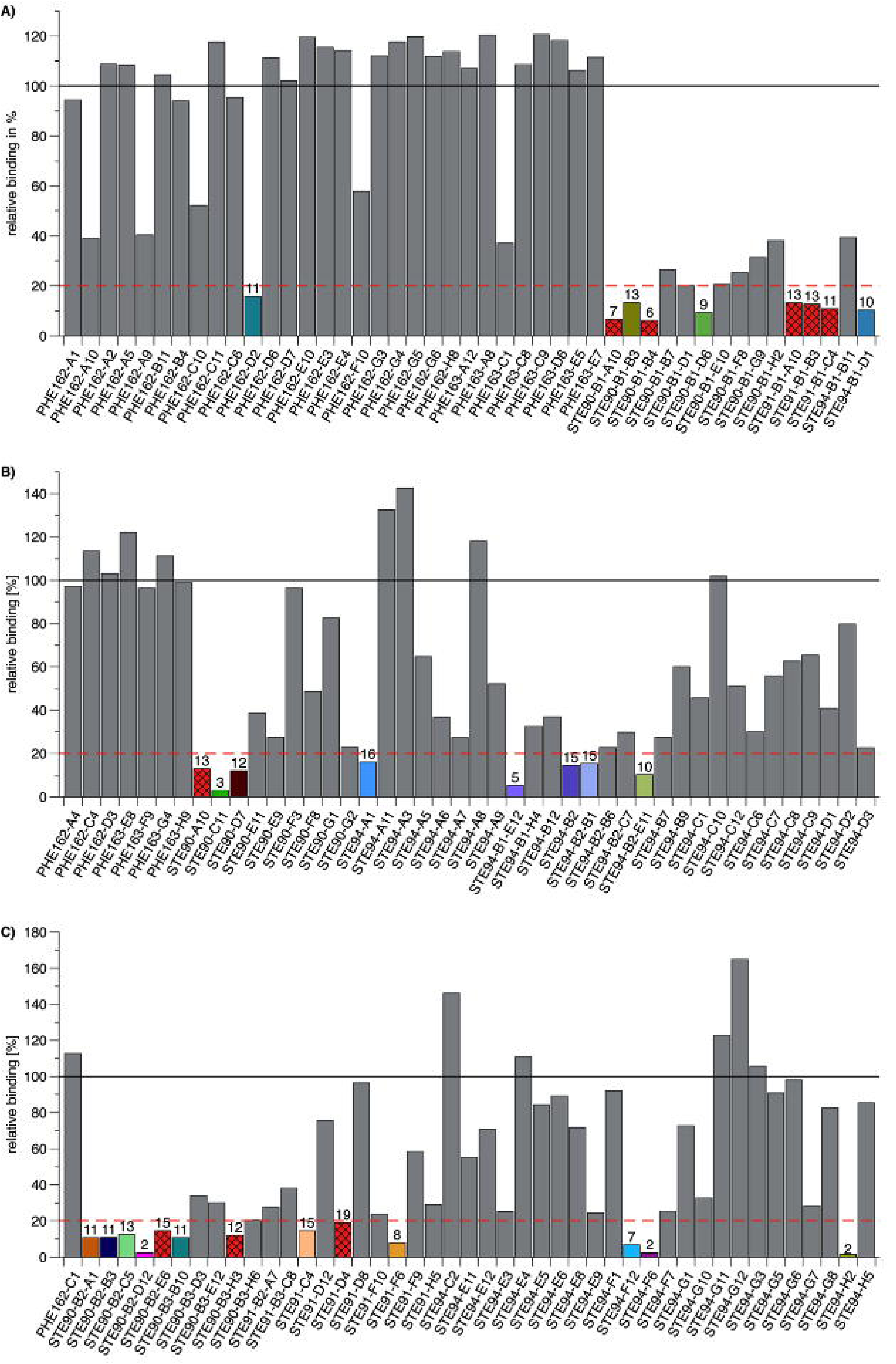
Inhibition of SARS-CoV-2 spike protein binding cells (flow cytometry). (A) Inhibition prescreen of 135 scFv-Fc antibodies on ACE2 positive cells using 1500 nM antibody and 50 nM spike protein (30:1 ratio). The antibodies selected for detailed analysis are marked in colors. Bars with red background and crossed-striped indicate scFv-Fcs not further analyzed. Data show single measurements.

**Table 1.** Overview on inhibiting antibodies

| Antibody name | VH | Germinality index VH [%] | VL | Germinality index VL [%] | EC50 ELISA [nM] |  |  | flow cytometry spike binding inhibition assay |  |  |  |
| --- | --- | --- | --- | --- | --- | --- | --- | --- | --- | --- | --- |
|  |  |  |  |  | RBD | S1 | S1-S2 | IC 50 [nM] with 50 nM spike | Molar ratio antibody arm: spike | IC 50 [nM] with 10 nM RBD | Molar ratio antibody arm: RBD |
| PHE162-D2 | IGHV4-31 | 98.9 | IGLV1-40 | 98.5 | 3.3 | 3.9 | 7.5 | 160 | 3.20 | 34.0 | 3.40 |
| STE90-A10 | IGHV3-66 | 98.6 | IGKV1-5 | 94.4 | / | / | / | / | / | / | / |
| STE90-C11 | IGHV3-66 | 98.6 | IGKV1-9 | 100 | 0.2 | 0.2 | 0.3 | 2.56* | 0.05* | 0.99* | 0.03* |
| STE90-D7 | IGHV3-66 | 98.6 | IGKV1-12 | 98.1 | n.d. | n.d. | n.d. | n.d. | n.d. | n.d. | n.d. |
| STE94-A1 | IGHV3-66 | 98.6 | IGLV1-40 | 100 | n.d. | n.d. | n.d. | 460 | 9.20 | 31.3 | 3.13 |
| STE90-B1-A10 | IGHV3-66 | 98.6 | IGKV1-5 | 98.1 | / | / | / | / | / | / | / |
| STE90-B1-B3 | IGHV3-66 | 98.6 | IGKV1-9 | 96.0 | / | / | / | 21.1 | 0.42 | 4.56 | 0.46 |
| STE90-B1-B4 | IGHV3-66 | 98.6 | IGKV1D-39 | 92.5 | / | / | / | / | / | / | / |
| STE90-B1-D6 | IGHV3-53 | 97.7 | IGKV1D-39 | 96.4 | / | / | / | 80.6 | 1.61 | 81.6 | 0.82 |
| STE90-B2-A1 | IGHV3-66 | 98.6 | IGKV1D-39 | 92.8 | 1.4 | 2.6 | 5.2 | 620 | 12.4 | 19.3 | 1.93 |
| STE90-B2-B3 | IGHV3-66 | 98.6 | IGKV1-5 | 100 | n.d. | n.d. | n.d. | n.d. | n.d. | n.d. | n.d. |
| STE90-B2-C5 | IGHV3-66 | 98.6 | IGKV1-5 | 100 | n.d. | n.d. | n.d. | 420 | 8.40 | 640 | 64.0 |
| STE90-B2-D12 | IGHV3-66 | 98.6 | IGKV1-9 | 100 | 0.3 | 0.4 | 0.4 | 27.1 | 0.54 | 4.49 | 0.45 |
| STE90-B2-E6 | IGHV3-53 | 100 | IGKV1D-39 | 98.1 | / | / | / | n.d. | n.d. | n.d. | n.d. |
| STE90-B3-B10 | IGHV3-66 | 98.6 | IGKV1-5 | 100 | n.d. | n.d. | n.d. | 250 | 5.0 | 490 | 49.0 |
| STE90-B3-H3 | IGHV3-66 | 98.6 | IGKV1D-12 | 100 | / | / | / | / | / | / | / |
| STE91-B1-A10 | IGHV3-66 | 98.6 | IGLV3-1 | 89.4 | / | / | / | / | / | / | / |
| STE91-B1-B3 | IGHV3-66 | 98.6 | IGLV3-21 | 100 | / | / | / | / | / | / | / |
| STE91-B1-C4 | IGHV3-66 | 98.6 | IGLV3-1 | 85.9 | / | / | / | / | / | / | / |
| STE91-C4 | IGHV3-66 | 98.6 | IGLV3-21 | 91.6 | 4 | 7.9 | n.d. | 36.4 | 0.73 | 7.15 | 0.72 |
| STE91-D4 | IGHV3-66 | 98.6 | IGLV3-1 | 89.4 | / | / | / | / | / | / | / |
| STE91-F6 | IGHV3-66 | 98.6 | IGLV1-40 | 100 | 2.3 | 3.8 | 5.9 | 30.0 | 0.60 | 5.99 | 0.60 |
| STE94-B1-D1 | IGHV3-66 | 98.6 | IGLV3-21 | 95.0 | 1 | 0.9 | 2.5 | 15.5 | 0.31 | 3.09 | 0.31 |
| STE94-B1-E12 | IGHV3-66 | 98.6 | IGLV3-1 | 85.0 | 0.4 | 0.7 | 1.6 | 11.5 | 0.23 | 2.49 | 0.25 |
| STE94-B2 | IGHV3-66 | 98.6 | IGLV3-21 | 98.3 | 5.1 | 4.8 | n.d. | n.d. | n.d. | 5.55 | 0.56 |
| STE94-B2-B1 | IGHV3-66 | 98.6 | IGLV1-40 | 100 | 1.2 | 2 | 7.5 | 140 | 2.80 | 3.35 | 0.34 |
| STE94-B2-E11 | IGHV3-66 | 98.6 | IGLV1-51 | 91.3 | 4.7 | 7.7 | n.d. | 39.1 | 0.78 | 6.25 | 0.63 |
| STE94-F6 | IGHV1-69 | 96.3 | IGKV1D-39 | 96.2 | 0.2 | 0.3 | 0.3 | 10.3 | 0.21 | 2.06 | 0.21 |
| STE94-F12 | IGHV3-66 | 98.6 | IGLV3-21 | 93.3 | 0.7 | 1.2 | 2.4 | 43.7 | 0.87 | 8.74 | 0.87 |
| STE94-H2 | IGHV3-66 | 98.6 | IGKV1-9 | 100 | 0.3 | 0.4 | 0.5 | 5.55 | 0.11 | 1.11 | 0.11 |

### Antigen binding and cell-based inhibition by IgG

The 30 antibodies were screened in a cytopathic effect (CPE)-based neutralization screening assay (data not shown) to select antibodies for further characterization as IgG. This assay was performed with 250 pfu/well SARS-CoV-2 and 1 μg/mL (∼10 nM) scFv-Fc. CPE is characterized by rounding and detachment clearly visible in phase contrast microscopy upon SARS-CoV-2 infection within 4 days, while uninfected cells maintained an undisturbed confluent monolayer. The best neutralizing 19 scFv-Fc were re-cloned and produced as human IgG in 50 mL culture scale, with yields ranging from 12 to 93 mg/L. EC_50_ values of binding to RBD, S1 or S1-S2 were determined (Figure 2) and are shown in Table 1. Antibodies STE90-C11, STE90-B2-D12, STE94-F6 and STE94-H2 showed EC_50_ values ranging between 0.2-0.5 nM on all antigens tested. Inhibition of ACE2 binding was assessed at concentrations from 100 nM - 0.3 nM IgG using 10 nM RBD (Supplementary Data 5A) or 500 nM - 1.5 nM IgG using 50 nM S1-S2 spike (Supplementary Data 5B) in the previously described cell-based inhibition assay (Bertoglio et al., 2021). The extent of inhibition by STE90-C11 of both RBD and S1-S2 was validated in a further cellular assay with ACE2 expressing cells (Supplementary Data 6). The best antibodies showed IC_50_ values of 2.6-15 nM IgG for 50 nM S1-S2 spike, respectively 1-3 nM IgG for 10 nM RBD (Table 1). For 50% inhibition the best molar ratios were from 0.02-0.3:1 (antibody binding site:antigen).

**Fig. 2.**
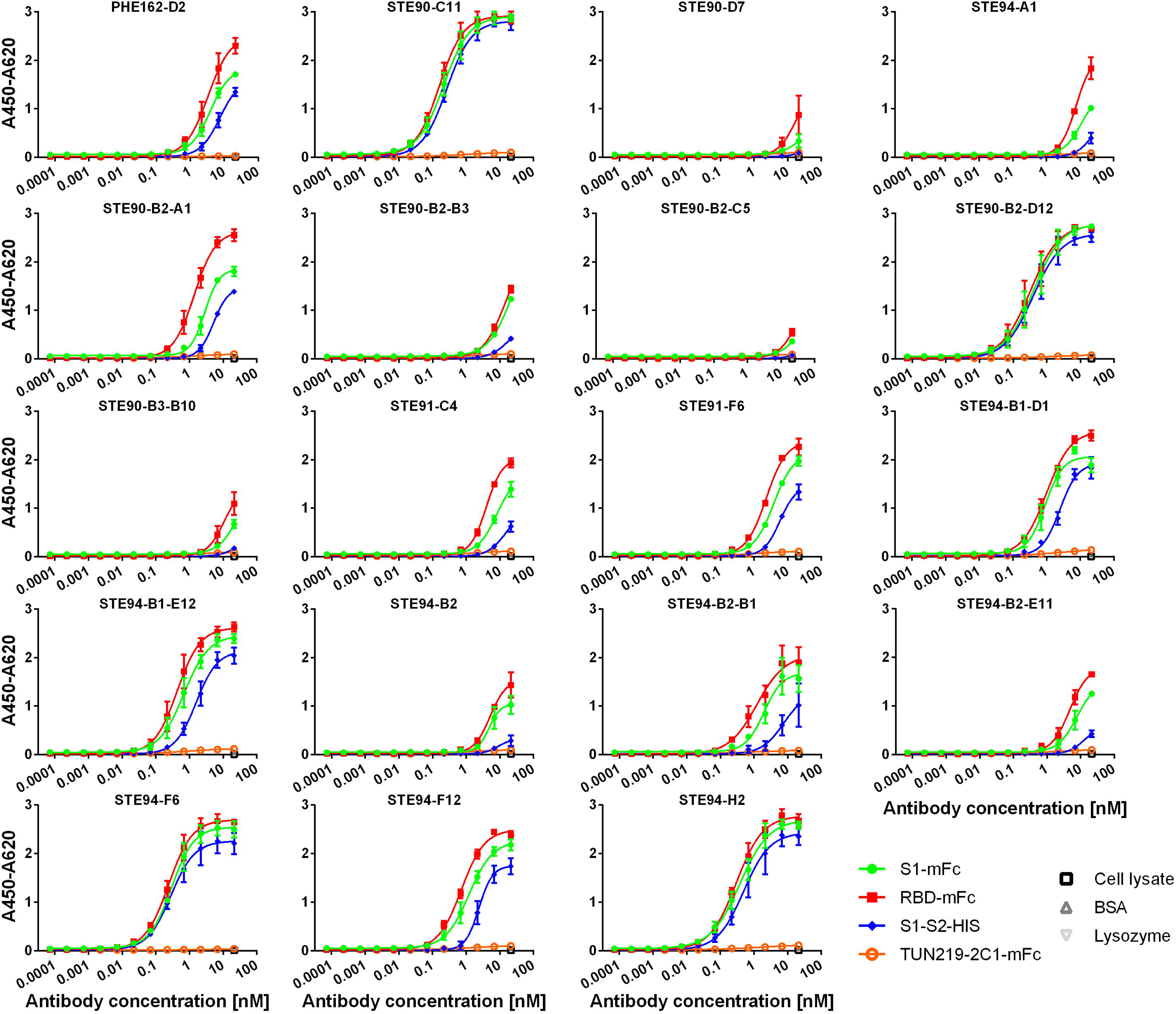
Determination of EC50 in ELISA. Binding in titration ELISA of the IgGs to (A) RBD (fusion protein with murine Fc part), (B) S1 (fusion protein with murine Fc part) or (C) S1-S2 (fusion protein with His tag). An unrelated antibody with murine Fc part (TUN219-2C1), human HEK293 cell lysate, BSA and lysozyme were used as controls. Experiments were performed in duplicate and mean ± s.e.m. are given. EC50 were calculated with GraphPad Prism Version 6.1, fitting to a four-parameter logistic curve.

### Antibodies from COVID19 patient libraries neutralize active SARS-CoV-2 virus

To analyze the neutralization capacity of the antibodies, the 19 IgGs were first screened in a plaque titration assay using active ∼15 pfu SARS-CoV-2 virus. The antibody Palivizumab was used as control antibody (Figure 3A). The antibody STE90-C11 was chosen to be further analyzed. To confirm its neutralizing activity, a plaque assay using ∼150 pfu was performed and the determined IC_50_ was 0.56 nM in the IgG format (Figure 3B).

**Fig. 3.**
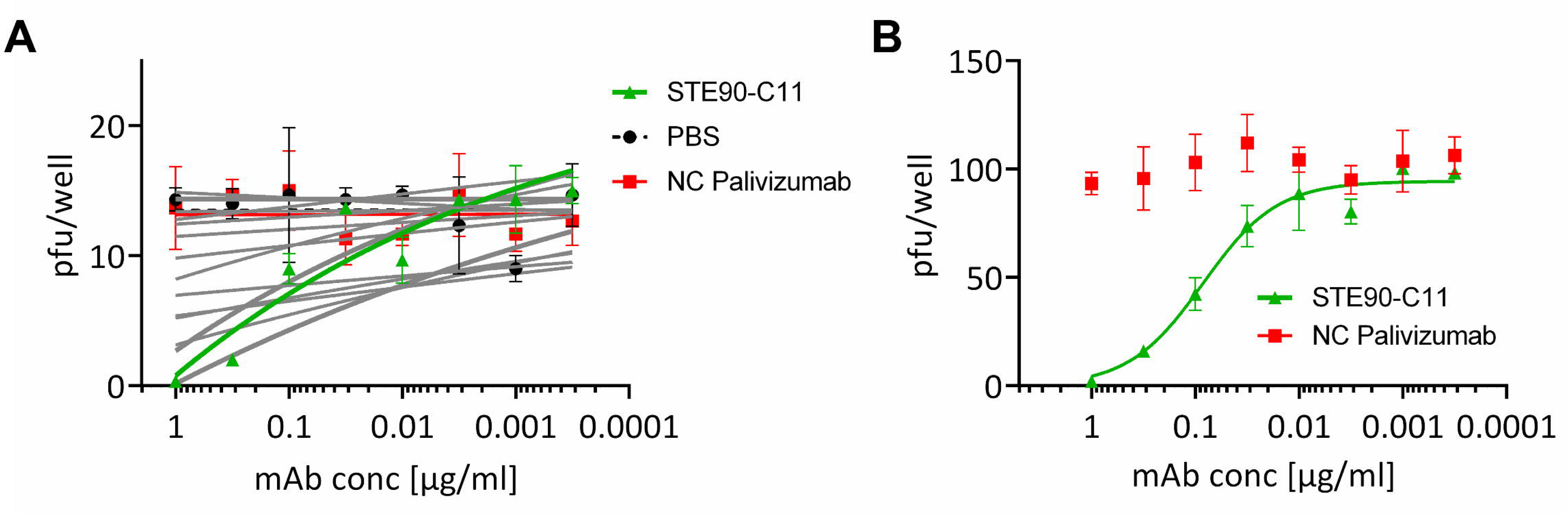
*In vitro* neutralization of authentic SARS-CoV-2. (A) Neutralization screening of ∼15 pfu SARS-CoV-2 by 19 anti-RBD SARS-CoV-2 IgGs. (B) STE90-C11 titration on ∼150 pfu SARS-CoV-2 to determine the IC50. Neutralization assays were performed in triplicates, mean ± s.e.m. are given. Palivizumab was used as isotype control in both assays. IC50 were calculated with OriginPro using the Logistic5 Fit.

### STE90-C11 is specific for SARS-CoV-2 and binds most RBD mutations including B.1.617

A detailed characterization of the best neutralizer was performed in respect to cross-reaction to other coronaviruses, binding to known RBD mutations, as well as biochemical and biophysical properties. The binding of STE90-C11 to other coronaviruses SARS-CoV-1, MERS-CoV, HCov-HKU1, HCoV-229E and HCoV-NL63 Spike proteins was tested by titration ELISA (Supplementary Data 7). STE90-C11 bound specifically SARS-CoV-2 and did not show any cross reactions to other human-pathogenic coronaviruses.

Because first SARS-CoV-2 mutants in patients are being identified (www.gisaid.org and (Shi et al., 2020)) and more mutations within the RBD have been characterized to arise under antibody selection pressure (Baum et al., 2020), S1 subunits harboring single point mutations described in GISAID in the RBD were produced and binding of STE90-C11 to those mutants was tested. In addition a variant with seven point mutations (V367F, N439K, G476, V483A, E484K, G485R and F486V) was analyzed (S1-7PM). The EC50 values of STE90-C11 are given in Figure 4 and the corresponding ELISAs are given in Supplementary Data 8, including further binding assays comparing STE90-C11 to REGN10933, REGN10987, CR3022 and CB6. All S1 or S1-S2 constructs were still able to bind recombinant ACE2 (Supplementary Data 9), indicating correct folding. The antibody STE90-C11 showed reduced binding to N501Y and lost binding to K417N/K417T but was able to bind to all other investigated RBD mutations including a S1-7PM mutant and more relevant to E484K, N439K and L452R which are the RBD mutations in the emerging B.1.525, B.1.526, B.1.1.33, B.1.258 and B.1.429/B.1.427 variants. Most important is the binding to B.1.617 (L452R+E484Q) which is currently emerging in India and other parts of the world. Subsequently, the inhibition of S1 binding to ACE2 for the RBD mutants bound best by STE90-C11 was confirmed in the cell based inhibition assays (Supplementary Data 10). The inhibition of B.1.617 was as slightly better than the inhibition of the wildtype.

**Fig. 4.**
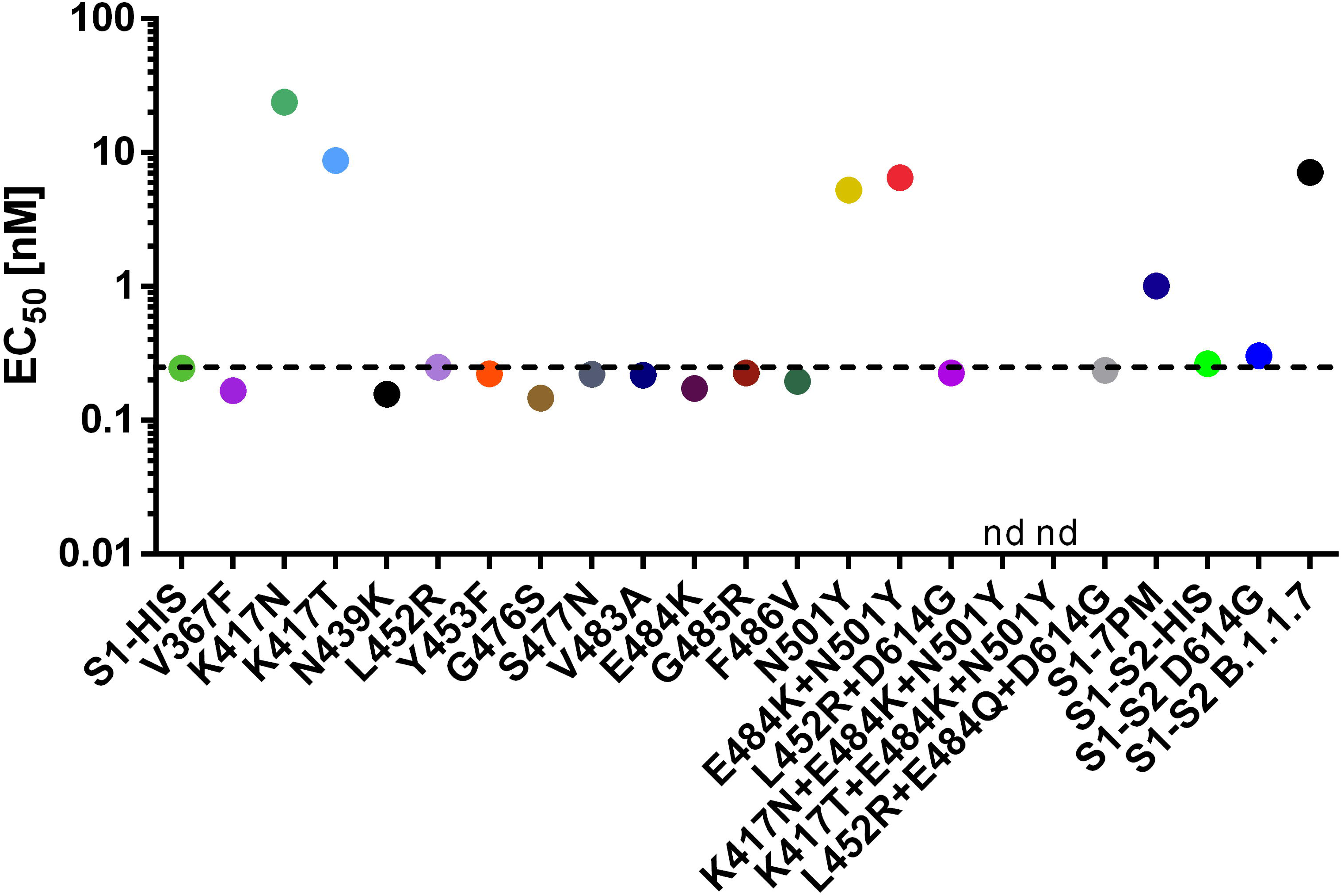
Cross reactivity analysis of STE90-C11. Binding to S1 or S1-S2 of different SARS-CoV-2 RBD mutations identified in virus variants from COVID-19 patients analyzed by ELISA. The EC50 values are given for each analyzed mutant. The corresponding ELISAs are given in Supplementary Data 8. n.d., not determinable. S1-7PM contains the seven mutations V367F, N439K, G476, V483A, E484K, G485R and F486V.

### STE90-C11 is protective in vivo in the hamster challenge model

The SARS-CoV-2 neutralizing effect of STE90-C11 was tested *in vivo* in a Syrian hamster challenge model. In this model, hamsters were intranasally infected with 1×10 ^5^ pfu genuine SARS-CoV-2 and 2 h later treated with 3.7 mg/kg or 37 mg/kg STE90-C11 or with PBS in the control group. The virus titer in the lung of treated animals showed a dose dependent reduction of viral load on day 3 and 5 after infection (Figure 5A). The measured mean viral load was 8.3×10^3^ pfu with 37 mg/kg IgG compared to 1.5×10^6^ pfu in the PBS control at day 3. The hamster model is further characterized by a rapid weight loss in the first days after SARS-CoV-2 infection but also rapid recovery of body weight from around one week post infection on. Animals treated with the higher STE90-C11 dose showed a reduced loss of weight and a faster weight recovery compared to the PBS control (Figure 5B).

**Fig. 5.**
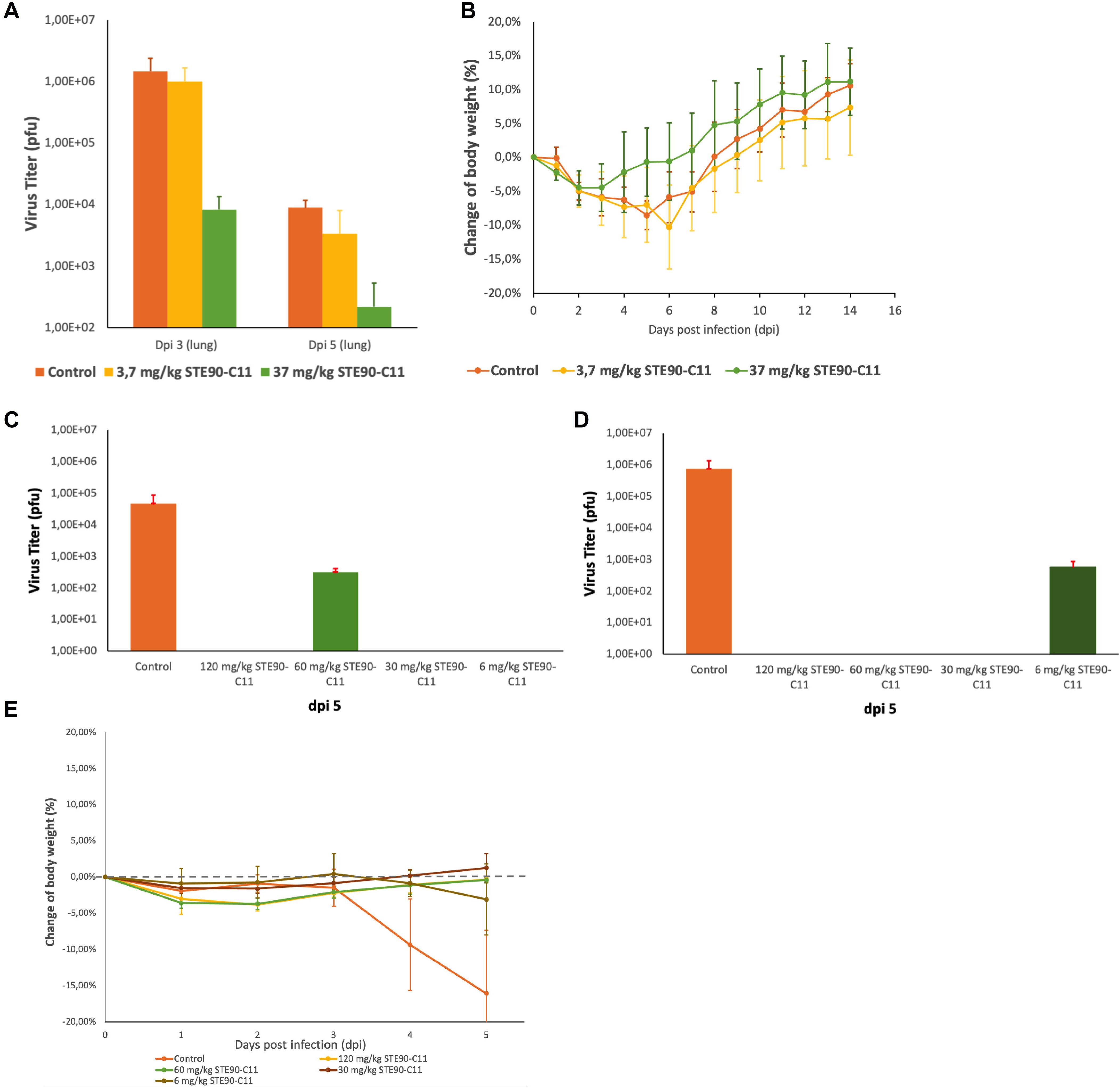
*In vivo* protection of STE90-C11 in a Syrian hamster challenge model and in the transgenic mice model. (A,B), Hamster model. Challenging of 9 animals per group with authentic SARS-CoV-2 and treatment with 3.7 mg/kg, 37 mg/kg STE90-C11 or with PBS. (A) Quantification of SARS-CoV-2 plaque-forming units (PFU) from lung homogenates after SARS-CoV-2 challenge 3 days after infection (3 Dpi) or 5 days (5 Dpi). (B) Body weight of hamsters after SARS-CoV-2 challenge from day 0 to 14. The mean values ± SEM from 9 animals per group are given. Mice model: 2-4 animals per group were treated with 6, 30, 60 or 120 mg/kg STE90-C11 or with PBS as control and 1 h later infected with authentic SARS-CoV-2. (C-E), K18hACE2 mouse model. (C, D), Quantification of SARS-CoV-2 plaque-forming units (PFU) from lung homogenates after SARS-CoV-2 challenge 5 days after infection (5 dpi). Two independent experiments are shown. Experiment 1 was performed with 1000 pfu and experiment 2 with 2000 pfu SARS-CoV-2. (E) Body weight of the mice after SARS-CoV-2 challenge from day 0 to 5 from three independent experiments. The mean values ± SEM from 2-4 animals per group are given.

### STE90-C11 is protective in vivo in the transgenic ACE2 mice challenge model

The SARS-CoV-2 neutralising effect of STE90-C11 was further tested in vivo in the transgenic human ACE2 mice model (K18hACE2). Mice were first treated with 6, 30, 60 or 120 mg/kg STE90-C11 or with PBS in the control group. After 1h, they were inoculated intranasally with 1,000 or 2,000 pfu genuine SARS-CoV-2, or with PBS in the control group. The virus titer was measured with 4.7×10^4^ pfu in the first experiment with 1,000 pfu and 7.4×10^5^ pfu in the experiment with 2,000 pfu. Only in the two experiments, one with 60 mg/kg and one with 6 mg/kg a remaining viral load of 5.6×10^2^, respectively 3.1×10^2^ pfu, was detected. In all other mice treated with STE90-C11, the virus was completely removed from the lungs (Figure 5C and 5D). In the control group, a significant weight loss was measured 4 days post infection which was not observed in the groups with STE90-C11 treatment (Figure 5 E).

### SARS-CoV-2 escape mutant screening leads to increased K417x but not N501x mutations

An *in vivo* escape mutant screening experiment was performed according to Baum et al. (Baum et al., 2020) to determine RBD mutants which are not neutralized by STE90-C11. Here, serial dilutions of STE90-C11, respectively Palivizumab as control antibody, were used to infect Calu-3 cells with active wt SARS-CoV-2. The supernatant with the propagated virus particles were used for the second round of neutralization and infection. After the second passage, the viruses were analyzed by DNA sequencing. In the experiment with STE90-C11 4.7% of the remaining viruses were wt, 81.9% had the K417T mutations, 3.35 K417T+A475V, 0.3% A475V, 0.3% N481H, 0.12% N481Y and 0.11% N501T. In the control experiment with a non related antibody and no selection pressure 89.9% of the remaining viruses were wt, 0.32% N481H, 0.14% K417T, 0.13% N501T and 0.1% Q414K. This experiment shows that the K417x mutation was not neutralized by STE90-C11, but no selection pressure was found on the N501x mutations, indicating a sufficient neutralization of these variants by STE90-C11.

### Biochemical characterization of STE90-C11

Aggregation propensity is an important quality parameter in therapeutic antibody development. STE90-C11 showed no relevant aggregation under normal conditions (pH7.4, 4°C in PBS), heat stress conditions (pH7.4, 45°C, 24h in PBS) and pH stress (pH3, 24h, RT) (Supplementary Data 11). A further test to exclude polyreactivity was performed on different antigens (DNA, LPS, lysozyme and EXPI cell lysate) with the FDA-approved antibody Avelumab as control (Supplementary Data 12). Here, STE90-C11 showed equal or less unspecificity compared to Avelumab. The monovalent affinity of STE90-C11 was determined by Bio-Layer Interferometry (BLI) in three different setups (Supplementary Data 13). In the first setup, RBD-mFc was immobilized using anti-Fc capture tips and the K_D_ of STE90-C11 in monovalent Fab format was determined as 8.1 nM. In the second setup, the IgG was immobilized using a Fab2G capture tip and the K_D_ of monovalent S1-His was determined to be 1.6 nM. In the third assay, the IgG was immobilized using a protein A capture tip, resulting in a measured K_D_ of 6.5 nM to S1-His, confirming the affinity of STE90-C11 is in the low nanomolar range. STE90-C11 did not bind to RBD on immunoblots after treatment at 95°C and SDS-gel separation under reducing conditions, while it was able to detect RBD when separated under non-reducing conditions heated either at 95°C or 56°C (Supplementary Data 14). These data suggest that the antibody recognizes a conformational epitope.

### STE90-C11 binds at the ACE2-RBD interface

To get further insight into the neutralizing mechanism of STE90-C11, a complex of STE90-C11 Fab and SARS-CoV-2 RBD (22 kDa fragment) was prepared and subjected to crystallization screening. X-ray diffraction images collected from resulting crystals yielded a dataset to an overall resolution limit of 2.0 Å (Supplementary Data 15). After solving the structure by molecular replacement a model was built into the electron density (Figure 6). Figure 6A shows the binding of STE90-C11 to RBD and Figure 6B the binding of ACE2 to RBD. STE90-C11 binds SARS-CoV2 RBD with an interface area of 1113 Å^2^ (Figure 6C, Supplementary Data 16). Roughly 60% of this area can be contributed to the VH segment forming up to ten hydrogen bonds at the same time and region dominated by hydrophobic interaction (V99, A100, Y33, Y52) (Figure 6D). The remaining 40% are provided by the VL segment contributing eight additional hydrogen bonds to stabilizing the interaction (Figure 6E). All six CDR loops contribute to the interaction between STE90-C11 Fab and SARS-CoV-2 RBD. The superposition of the RBDs of the STE90-C11:SARS-CoV-2 RBD complex with a ACE2:SARS-CoV-2 RBD complex (Lan et al., 2020) resulted in a low Cα root mean square distance of 0.532 Å (183 atoms) indicating that binding of STE90-C11 did not induce substantial changes in the RBD. The superposition further revealed that the neutralization mechanism of STE90-C11 is based on directly competing for the ACE2 binding side, as the interaction interface on the RBD of both molecules almost completely overlap. Further superposition of the STE90-C11:SARS-CoV-2 RBD complex model onto cryo-EM models of the whole SARS-CoV-2 spike protein (Walls et al., 2020) indicate that due to steric hindrance STE90-C11 like ACE2 is expected to only bind to the open conformation of the spike protein (Supplementary Data 17).

**Fig. 6.**
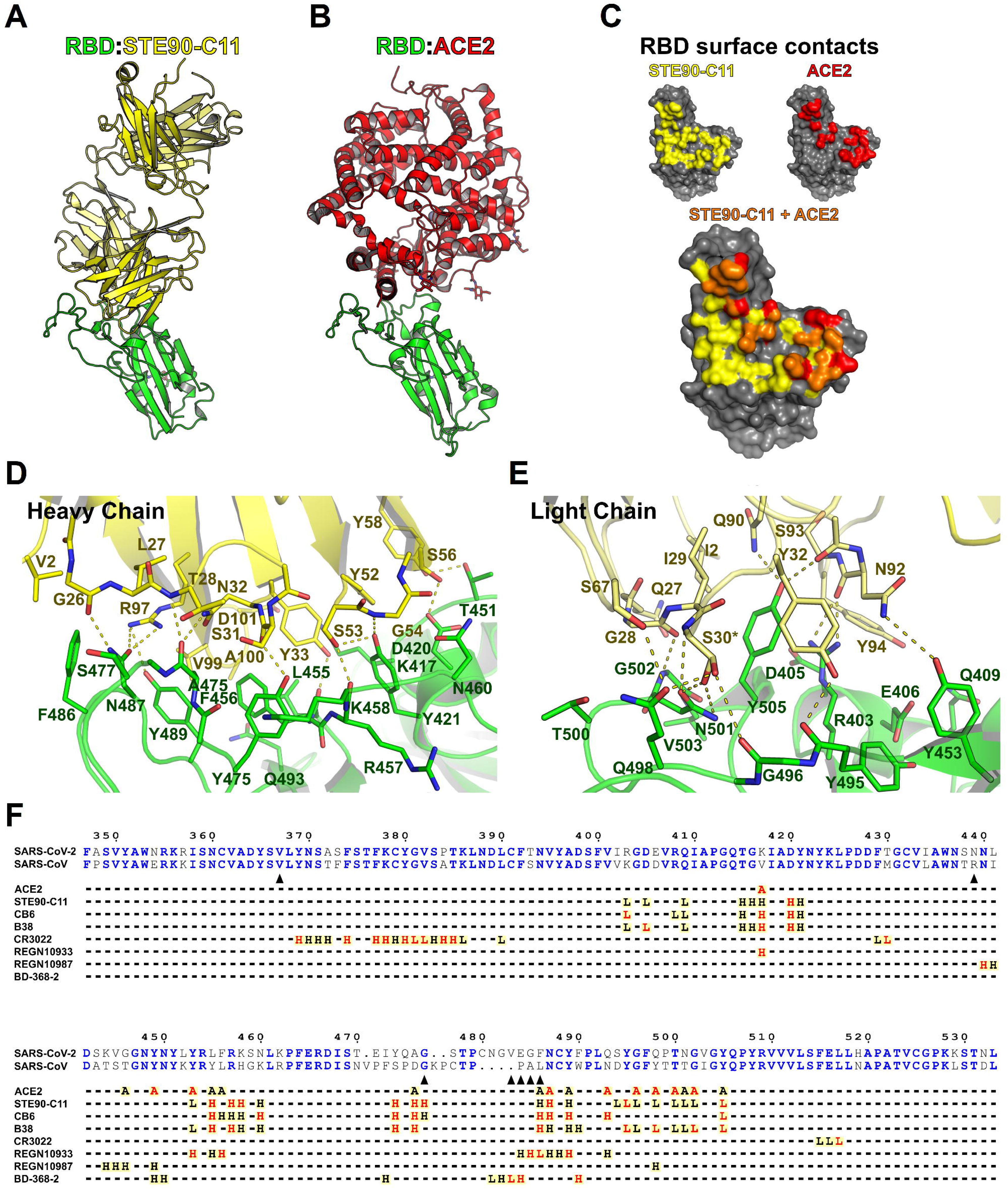
The crystal structure of STE90-C11 in complex with SARS-CoV-2-RBD. (A+B) The structure of the complex between RBD (green) and STE90-C11 (A; heavy chain: Yellow; light chain: pale yellow) or ACE2 (B; red; PDB: 6M0J (Lan et al., 2020)) depicted in cartoon representation after superposition of the RBD. (C) Surface representation of the RBD with the binding surface (cutoff 4Å) of STE90:C11 colored in yellow and of ACE2 colored in red. The competitive binding surface is colored in orange. (D+E) Detailed view of the interactions between RBD and the heavy chain (D) and light chain (E) of STE90-C11. Possible hydrogen bonds are displayed as dashed yellow lines and residues with alternate conformations are marked with an asterisk. (F) Sequence alignment of RBD. Depicted is a part of the sequence alignment of the SARS-CoV-2 and SARS-CoV S-protein (UniProt: P0DTC2; P59594). Identical residues are depicted as bold blue letters. Under the alignment contacts between residues of the S-protein and ACE2 (PDB: 6M0J (Lan et al., 2020)), STE90-C11 (PDB: 7B3O), CB6 (PDB: 7C01 (Shi et al., 2020)), B38 (PDB: 7BZ5 (Wu et al., 2020)) CR3022 (PDB: 6W41 (Yuan et al., 2020)), REGN10933, REGN10987 (PDB: 6XDG (Hansen et al., 2020)) and BD-368-2 (PDB: 7CHF (Cao et al., 2020)) are marked by an A (ACE2), H (heavy chain) or L (light chain). A black letter indicates a distance under 4Å and a red letter under 3.2Å between non-hydrogen atoms of the RBD and the respective protein. Black arrows indicated residues often exchanged in the RBD of SARS-CoV-2, which were tested for their influence on STE90-C11 binding in this work. Distances were calculated with CNS (Brünger et al., 1998) and the representation was prepared utilizing ESPript (Robert and Gouet, 2014).

## Discussion

For the treatment of COVID-19 but also to protect risk groups, human anti-SARS-CoV-2 antibodies are a promising therapeutic option. Human recombinant antibodies were successfully used for the treatment of other viral diseases. The antibody mAb114 (Ansuvimab-zykl) (Corti et al., 2016) and three-antibody cocktail REGN-EB3 (Atoltivimab/maftivimab/odesivimab) (Pascal et al., 2018) were FDA approved in 2020 and showed a good efficiency in clinical trials against Ebola virus especially in comparison to Remdesivir (Mulangu et al., 2019). For the treatment of a severe respiratory infection of infants caused by the respiratory syncytial virus (RSV), the antibody Palivizumab is approved by EMA and FDA (van Mechelen et al., 2016; Subramanian et al., 1998). These anti-viral antibodies can be used as blueprint to develop therapeutic antibodies against SARS-CoV-2. Currently, monoclonal antibodies against SARS-CoV-2 were selected by rescreening memory B-cells from a SARS patient (Pinto et al., 2020), selected from COVID-19 patients by single B-cell PCR (Cao et al., 2020; Shi et al., 2020) or FACS sorting (Kreer et al., 2020), selected from transgenic mice followed and from patients by single B-cells FACS sorting (Hansen et al., 2020) or using phage display using universal or patient libraries including different antibody formats (Bertoglio et al., 2021; Chi et al., 2020; Li et al., 2020a; Liu et al., 2020; Ma et al., 2021; Noy-Porat et al., 2020; Parray et al., 2020; Zeng et al., 2020).

In this work, antibody phage display immune libraries were constructed using lymphocytes from six local convalescent COVID-19 patients. 197 unique antibodies were selected against RBD. We intently focused on RBD aiming to select antibodies which directly interfere with the interaction between spike and ACE2, to avoid potential antibody dependent enhancement (ADE), especially in patients with severe symptoms. ADE during coronavirus entry has been described for both MERS (Wan et al., 2020) or SARS (Wang et al., 2014). It is defined as “enhancement of disease severity in an infected person or animal when an antibody against a pathogen…worsens its virulence by a mechanism that is shown to be antibody-dependent” (Arvin et al., 2020). Furthermore, immune dysregulation and lung inflammation was also caused by anti-spike antibodies in an acute SARS-CoV infection (Liu et al., 2019). On the other hand, Quinlan *et al*. (Quinlan et al., 2020) reported that animals immunized with RBD SARS-CoV-2 did not mediate ADE. A therapeutic antibody directed against RBD and the usage of a Fc-part which is not binding to Fc-gamma receptors (“silent Fc”) would reduce the risk of ADE, in particular because ADE cannot be fully predicted from *in vitro* studies or from animal models (Arvin et al., 2020). Winkler and et al. argue, that an intact Fc part is needed for optimal therapeutic protection (Winkler et al., 2021). The ACTIV-3/TICO LY-CoV555 Study Group describes no efficacy of LY-COV-555 in hospitalized patients and discussed also ADE as potential explanation for the lacking efficacy (ACTIV-3/TICO LY-CoV555 Study Group et al., 2021). Because, the role of ADE is not fully deciphered in case of SARS-CoV-2 (Lee et al., 2020) and LY-CoV555 was not effective in patients with severe symptoms, it is advised to use a silent Fc part to address hospitalized patients with moderate to severe symptoms. Therefore, all assays with STE90-C11 IgGs were performed with a silenced Fc part.

The selected anti-RBD antibodies were analyzed for RBD::ACE2 inhibition in the scFv-Fc format. The advantage of the IgG like bivalent scFv-Fc format is that it speeds up analysis by requiring only one cloning step and provides high expression yields in small scale format (Bertoglio et al., 2021; Jäger et al., 2013; Wenzel et al., 2020a). 30 scFv-Fc antibodies inhibited the binding of the spike protein to ACE2-expressing living cells. Interestingly, the majority of the VH V-genes of the selected anti-RBD antibodies and 28 of these 30 inhibiting antibodies is the VH3-66 V-gene showing strong enrichment of this V-gene in the immune response of COVID-19 patients against RBD. This is in accordance with Cao *et al*. (Cao et al., 2020) who found that enriched use of VH3-53 or VH3-66. VH3-53 and VH3-66 V-genes are closely related. Robbiani *et al*. (Robbiani et al., 2020) showed an enrichment of VH3-30, VH3-53 and VH3-66 and Hansen et al. (Hansen et al., 2020) reported a bias towards VH3-53. The VH3-53 and VH3-66 V-genes are closely related. In contrast, Ju et al. (Ju et al., 2020) identified mainly VH1-2, VH3-48 and VH3-9 V-genes for their selected anti-RBD antibodies from one patient and Brouwer et al. (Brouwer et al., 2020) identified a strong enrichment of VH1-69, VH3-30 and VH1-24 in three patients, but also an enrichment of VH3-53 and VH3-66. VH3-66 antibodies against RBD were also selected from the naive HAL9/10 antibody gene libraries made long before the SARS-CoV-2 outbreak (Kügler et al., 2015) but not significantly enriched (Bertoglio et al., 2021). The selected inhibiting antibodies in our study (Table 1) all have a high germinality index and are therefore very similar to the not hypermutated human germline, promising low immunogenicity when used as a therapeutic agent. Only some light chains showed a major difference to their closest related V-gene. Kreye *et al*. (Kreye et al., 2020) also described that antibodies they selected from COVID-19 patients were very close to the germline genes. Interestingly, the VH V-genes of these immune libraries are closer to their germline compared to the anti-RBD antibodies selected from the naive HAL9/10 antibody gene libraries (Bertoglio et al., 2021).

In the cell-based inhibition assay, some of the antibodies showed a molar antibody:spike monomer ratio lower then 1:1. Similar behaviour was also found by Bertoglio *et al*. (Bertoglio et al., 2020) and may be explained by the observation that three RBDs on the same spike trimer can be in different conformations (“up” and “down”), while only the RBDs in “up” positions are able to bind ACE2 (Walls et al., 2020) and therefore, the “down” RBDs are not fully accessible.

The monovalent affinity of STE90-C11 was measured with 1.6 nM-8.1 nM depending on the BLI setup. This is in the same range as BD-368-2 with 0.82 nM affinity to RBD (Cao et al., 2020), 3.3 nM for REGN10933 in the setup with the monovalent RBD (Hansen et al., 2020). 70 nM for B38 (Wu et al., 2020) or 0.8-4.7 nM for CB6 (Shi et al., 2020).

The analysis of the binding STE90-C11 to other human coronaviruses showed specificity for SARS-CoV-2. Comparison between the epitope of STE90-C11 and other published antibodies (Figure 5F) revealed that it binds in the same region as CB6 (Shi et al., 2020) and B38 (Wu et al., 2020). The selectivity of STE90-C11 and CB6 for SARS-CoV-2 over SARS-CoV RBD can mainly be attributed to the interaction with residues 473-476, as they are not involved in ACE2 binding and SARS-CoV RBD folds into a different conformation of this loop causing steric hindrance for antibody binding (Lan et al., 2020).

In addition to the *in vitro* neutralization of authentic SARS-CoV-2, the *in vivo* efficacy of STE90-C11 was demonstrated in the Syrian Hamster Challenge model (Kreye et al., 2020) and in the model using transgenic mice with human ACE2 receptor (Wu et al., 2020) in a dose dependent manner.

The binding of STE90-C11 to S1 containing RBD mutations observed in strains from COVID-19 patients was analyzed, showing a binding to most variants. As the cell-based inhibition analysis demonstrate, STE90-C11 is able to inhibit most analyzed mutants, validating a tolerance for RBD mutations. REGN10933 (Hansen et al., 2020), which is also binding at the RBD-ACE2 interface, showed a loss of neutralization in an assay using pseudoviral particles for the F486V and a reduced neutralization for both G485D and E484K mutations (Baum et al., 2020). Loss of binding to F486A is also described for VH-Fc ab8 (Li et al., 2020b). In the epitope analysis, REGN10933 has more molecular interactions within the region aa483-aa486 compared to STE90-C11. Currently, the variant B.1.1.7 is widely spread and STE90-C11 has reduced binding to N501Y and lost binding to mutants with the K417N/T mutations in combination with N501Y which occur in B.1.351 and B.1.1.28.1 variants. The structure suggest that the exchange N501Y may push the light chain CDR1-loop of the antibody further away while the exchange of K417N/T leads to an abolishment of a salt bridge to D101 of the heavy chain, similar to observations done for the COVOX-269 and COVOX-222, respectively (Dejnirattisai et al., 2021; Supasa et al., 2021). In COVOX-269 binding to the RBD-N501Y leads to a conformational change of light chain CDR3 loop around Y94. In the escape mutant experiment, no N501x mutants were enriched, indicating a sufficient neutralization of the N501x but not K417x mutation.

On the other hand, STE90-C11 binds strongly to the RBD mutations in the emerging SARS-CoV-2 variants B.1.429/B.1.427 (L452R), B.1.526 (E484K or S477N), B1.258Δ (N439K), B.1.525 (E484K), B.1.1.28.2 (E484K), B.1.1.33 (E484K) and B1.617 (L452R, E484Q). These variants are emerging, e.g. the frequency of B.1.429+B.1.427 reached 40% in January 2021 (Zhang et al., 2021) or B.1.258 with 59% in the samples sequenced in Czech in the last three month of 2020 (Brejová et al., 2021). B1.617 (L452R, E484Q) and derivates have superseded B.1.1.7 in India and the prevalence of this variant was majority of all sequenced viruses in End of April 2021 (https://outbreak.info/location-reports?loc=IND). We hypothesize that STE90-C11 is still binding to most analyzed RDB mutants because of both the wide interface area of 1133 Å^2^ and the extensive light chain contacts which may compensate the exchange of several amino acids. The calculated interface area of STE90-C11 VL is more than twice the area of REGN10933 and REGN10987 and slightly larger compared to CB6 or CR3022 (Supplementary Data 16).

In summary, the patient-derived SARS-CoV-2 neutralizing anti-RBD antibody STE90-C11 binding to the RBD-ACE2 interface maintains high similarity to the human germline V-genes VH3-66, the same family of many isolated anti-SARS-CoV-2 neutralizing antibodies. STE90-C11 is tolerant to most known RBD mutants especially those of the mutants B.1.429/B.1.427, B.1.526, B1.258Δ, B.1.535, B.1.617 and B.1.1.33 which are currently emerging. A phase Ib/II clinical trial with a full human IgG1 variant with FcγR silenced Fc STE90-C11 (COR-101) was started in April 2021 (ClinicalTrials.gov ID: NCT04674566) to assess safety and efficacy of COR-101 in hospitalized patients with moderate to severe COVID-19.

## STAR Methods

### Key resources table

| Reagent or Resource | Source | Identifier |
| --- | --- | --- |
| Phage Display Library |  |  |
| COVID-19 immune libraries | This paper | N/A |
| Antibodies |  |  |
| goat-anti-hlgG(Fc)-HRP | Sigma | Cat#A0170 |
| murine mAb 9E10 | Expressed for this paper in-house, original (Chan et al., 1987) | N/A |
| goat-anti-mIgG(Fc)-HRP | Sigma | Cat#A0168 |
| Hyper-Myc (TUN219-2C1-mFc) | Abcalis | Cat#ABC0503 |
| Avelumab | Expressed for this paper in-house, original (Boyerinas et al., 2015) | N/A |
| Palivizumab | Expressed for this paper in-house, original (Subramanian et al., 1998) | N/A |
| STE90-C11 | This paper | N/A |
| mouse anti-His epitope tag | Dianova | Cat#DIA-900-200 |
| mounse anti-penta His (APC conjugated) | Qiagen | Cat#3466 |
| goat anti-hlgG(Fc)-AP | Jackson ImmunoResearch | Cat#AB_2337599 |
| goat anti-mIgG-AP | Dianova | Cat#115-055-071 |
| mouse α-CD138 (FITC conjugated) | BioLegend | Cat#352304 |
| Bacterial and Virus strains |  |  |
| *E. coli* XL1 Blue MRF' | Stratagene | Cat#200230 |
| *E. coli* TG1 | Lucigen | Cat#605021 |
| SARS-CoV-2/Münster/FI110320/1/2020 | University of Münster, Germany | N/A |
| SARS-CoV-2 BetaCoV/Germany/BavPat1/2020 | Charité, Germany | N/A |
| Hyperphage | Produced in-house | N/A |
| M13KO7 helperphage | Produced in-house | N/A |
| Experimental Model: Cell lines |  |  |
| Expi293F | Thermo Fisher Scientific | Cat#A14527 |
| VeroE6 | ATCC | ATCC CRL-1586 |
| High Five | Thermo Fisher Scientific | Cat#B85502 |
| Biological Samples |  |  |
| COVID-19 patient samples | local hospitals | N/A |
| Healthy adult control samples (from pre-COVID-19 time) | LADR GmbH<br>Medizinisches Versorgungszentrum<br>Braunschweig | N/A |
| Chemicals, Peptides, and Recombinant Proteins |  |  |
| Spike protein (RBD; S1; S1-S2; N) | This paper | GenBank: MN908947 |
| ACE2 | This paper | GenBank: NM021804.3 |
| PEI 40 kDa | Polysciences | Cat#24765-1 |
| Ficoll | GE Healthcare | Cat#45-001-749 |
| TRIzol LS | Life Technologies | Cat#10296028 |
| Superscript IV | Thermo Fisher Scientific | Cat#18090010 |
| Trypsin | Sigma | Cat#9002-07-7 |
| IPTG (50 μM) | Sigma | Cat#367-93-1 |
| Gibco FreeStyle F17 expression media | Thermo Fisher Scientific | Cat#10388033 |
| HyClone SFM4Transfx-293 media | GE Healthcare | Cat#10015813 |
| HyClone Boost 6 supplement | GE Healthcare | Cat#11576341 |
| Laemmli sample buffer | Laemmli, 1970 | N/A |
| 5-Brom-4-Chlor-3-Indolyl-Phosphat/<br>Nitro Tetrazolium Blue Chloride | Thermo Fisher Scientific | Cat#B6149 |
| CHO CD efficient feed A | Thermo Fisher Scientific | Cat#A1023401 |
| ExCell 405 serum-free medium | Sigma | Cat#14405C |
| 2,3-(R,R)-butandiol | Alfa Aesar | Cat#513-85-9 |
| Critical Commercial Kits |  |  |
| Direct-zol RNA Miniprep Plus Kit | Zymo Research | Cat#R2072 |
| NucleoBond Xtra Midi Kit | Macherey-Nagel | Cat#12798402 |
| NucleoSpin Gel and PCR Clean-up Kit | Macherey-Nagel | Cat#11992242 |
| NucleoSpin Plasmid EasyPure | Macherey-Nagel | Cat#15162537 |
| NEB Luna Universal Probe One-Step RT-qPCR kit | NEB | Cat#E3006S |
| Deposited Data |  |  |
| Crystal structure of STE90-C11 in complex with SARS-CoV-2-RBD | This paper | PDB 7B3O |
| Oligonucleotides |  |  |
| Primer set for VH and VL amplification | (Wenzel et al., 2020b) | N/A |
| Recombinant DNA |  |  |
| Spike protein (RBD; S1; S1-S2; N) | Thermo Fisher/Gene Art | GenBank: MN908947 |
| ACE2 | Thermo Fisher/Gene Art | GenBank: NM021804.3 |
| Software and Algorithms |  |  |
| OriginPro (2019) | OriginLab | N/A |
| GraphPad Prism 6.1 | GraphPad | www.graphpad.com |
| VBASE | MRC Centre for Protein Engineering | www2.mrc-lmb.cam.ac.uk/vbase/ |
| VBASE2 | (Mollova et al., 2010) | www.vbase2.org |
| DataGraph (4.5.1) | DataGraph | www.visualdatatools.com/DataGraph/ |
| IncuCyte S3 GUI | Sartorius | N/A |
| Phenix | (Liebschner et al., 2019) | www.phenix-online.org |
| BD FACS Chorus | BD | N/A |
| PyMol | Schrödinger LLC | www.pymol.org/2/ |
| PISA | European Bioinformatics Institute | http://www.ebi.ac.uk/pdbe/prot_int/pistart.html |
| Octet qKe Data Analysis HT 11.0 | Fortebio/Sartorius | N/A |
| Other |  |  |
| Äkta Go/Äkta Pure | Cytiva | N/A |
| Profinia System | BIO-RAD | N/A |
| Precision XS microplate sample processor | BioTek | N/A |
| Sartorius IncuCyte S3 | Sartorius | N/A |
| BD FACSMelody Cell Sorter | BD | N/A |
| Octet qKe | Fortebio/Sartorius | N/A |
| PETRA III | DESY Hamburg | N/A |

#### Resource Availability

Further information and requests for resources and reagents should be directed to and will be fulfilled by the Lead Contact, Michael Hust

#### Materials Availability

All requests for resources and reagents should be directed to the Lead Contact author. This includes antibodies, plasmids and proteins. All reagents will be made available on request after completion of a Material Transfer Agreement.

#### Data and Code Availability

The structure data and the antibody sequence is available at PDB 7B3O.

### Experimental models and subject detail

#### SARS-CoV-2-infected individuals and sample collection

For the generation of human immune antibody libraries against SARS-CoV-2, blood from COVID-19 convalescent patients were collected from local hospitals. This was performed in accordance with the Declaration of Helsinki. All the voluntary donors were informed about the project and gave their consent. The use of blood samples for the development of antibody phage display libraries was approved by the ethical committee of the Technische Universität Braunschweig (Ethik-Kommission der Fakultät 2 der TU Braunschweig, approval number FV-2020-02). The donors have given their consent for study publication. A control serum was obtained from LADR Braunschweig.

### Method Details

#### Production of antigens in insect cells

The antigens were produced and purified as described before (Bertoglio et al., 2021; Korn et al., 2020). In brief, different domains or subunits of the Spike protein (GenBank: MN908947) were produced Baculovirus-free in High Five cells (Thermo Fisher Scientific, Schwerte, Germany) by transient transfection. High Five cells were cultivated at 27°C, 110 rpm in EX-CELL 405 media (Sigma Aldrich, Munich, Germany) and kept at a cell density between 0.3 – 5.5 ×10^6^ cells/mL. For transfection cells were centrifuged and resuspended in fresh media to a density of 4×10^6^ cells/mL and transfected with 4 µg plasmid/mL and 16 µg/mL of PEI 40 kDa (Polysciences). After 4 h to 24 h after transfection cells were diluted to a final density of 1x 10^6^ cells/mL. At 48 h after transfection, the culture volume was doubled. Cell supernatant was harvested five days after transfection in a two-step centrifugation (4 min at 180xg and 20 min at above 3500xg) and 0.2 µm filtered for purification.

#### Protein purification

Protein purification was performed as described before (Bertoglio et al., 2021) depending on the production scale in either 24 well filter plate with 0.5 mL resin (10 mL scale) or 1 mL column on Äkta go (Cytiva), Äkta Pure (Cytiva) or Profinia System (BIO-RAD). MabSelect SuRe or HiTrap Fibro PrismA (Cytiva) was used as resin for Protein A purification. For His-tag purification of Expi293F supernatant HisTrap FF Crude column (Cytiva) and for His-tag purification of insect cell supernatant HisTrap excel column (Cytiva) was used. All purifications were performed according to the manufacturer’s manual. Indicated antigens were further purified by size exclusion chromatography by a 16/600 Superdex 200 kDa pg (Cytiva). All antigens, antibodies and scFv-Fc were run on Superdex 200 Increase 10/300GL (Cytiva) on Äkta or HPLC (Techlab) on an AdvanceBio SEC 300Å 2.7 µm, 7.8×300 mm (Agilent) for quality control.

#### Serum ELISA

For COVID-19 convalescent serum analysis, sera were titrated in 11 steps (dilution ratio 1:√3) on 100 ng/well of RBD-His. As unspecificity controls, sera were tested on 100 ng/well BSA. All immobilization steps were performed in 0.05 M carbonate buffer (pH 9.6). Serum IgGs were detected using goat-anti-hIgG(Fc)-HRP (1:70,000, A0170, Sigma). Titration assays were performed in 96 well microtiter plates (Costar).

#### COVID-19 convalescent patient library construction

For the generation of human immune antibody libraries against SARS-CoV-2, blood from 6 donors showing a good antibody titer against RBD was used. The library construction was performed as described previously with minor modifications (Kügler et al., 2018). In brief, the peripheral blood mononuclear cells (PBMC) were extracted from the blood via Ficoll (GE Healthcare, Freiburg, Germany) and RNA isolated with TRIzol LS reagent (Life Technologies, Carlsbad, USA) and Direct-zol RNA Miniprep Plus kit (Zymo Research, Freiburg, Germany). For the plasma B-cell sorted library, plasma B-cells were double-stained with mouse α-CD19 APC-conjugated (MHCD1905, Thermo Fisher Scientific, Schwerte, Germany) and mouse α-CD138 (FITC conjugated) antibody (DL-101, BioLegend, San Diego, USA), sorted via FACSMelody (BD, Franklin Lakes, USA). Sorted cells were directly collected in TRIzol LS reagent and the RNA was extracted as indicated above. RNA was converted into cDNA using Superscript IV (Thermo Fisher Scientific, Waltham, USA) according to the manufacturer’s instructions. Library construction was achieved as described previously (Wenzel et al., 2020a) with slight modifications. In brief, specific primers for the VH chain, kappa light chain and lambda light chain were used to amplify antibody genes from the cDNA. These resulting PCR products were purified and again amplified with primers adding specific restriction sites for further cloning into the *E. coli* expression vector pHAL52. The vector pHAL52 is derived from the vector pHAL30 (Kügler et al., 2015) with an *AscI* restriction site in the VL stuffer and a *Sal*I restriction site in the VH stuffer for removal of uncut vector backbone. First, the light chain was cloned between the restriction sites *MluI-HF* and *NotI-HF*. In a second step the VH chain was cloned with the restriction sites *HindIII-HF* and *NcoI-HF* into previously cloned pHAL52-VL, additionally digested with *AscI*. The efficiency of library cloning was tested by colony PCR and the rate of complete scFv insertion was determined. The libraries were packaged with Hyperphage (Rondot et al., 2001; Soltes et al., 2007). Antibody phage were precipitated with PEG-NaCl and resuspended in phage dilution buffer (10 mM TrisHCl pH7,5, 20 mM NaCl, 2 mM EDTA). Resulting phage titer was determined by infection of *E. coli* XL1 blue MRF’.

#### Antibody selection using phage display

Antibody selection was performed as described previously with modifications (Russo et al., 2018). In brief, 5 µg of of S1-S2-His or RBD-His (produced in High Five cells) was diluted in carbonate buffer (50 mM NaHCO_3_/Na_2_CO_3_, pH 9.6) and coated onto the wells of a High binding 96 well microtiter plate (High Binding, Costar) at 4 °C overnight. Next, the wells were blocked with 350 µL 2% MBPST (2% (w/v) milk powder in PBS; 0.05% Tween20) for 1 h at RT and then washed 3 times with PBST (PBS; 0.05% Tween20). Before adding the libraries to the coated wells, the libraries (5×10^10^ phage particles) were preincubated with 5 µg of an unrelated scFv-Fc and 2% MPBST on blocked wells for 1 h at RT, to deprive libraries of human Fc fragment binders. The libraries were transferred to the antigen coated wells, incubated for 2 h at RT. After 10 washes, bound phage were eluted with 150 µL trypsin (10 µg/mL) at 37°C, 30 minutes and used for the next panning round. The eluted phage solution was transferred to a 96 deep well plate (Greiner Bio-One, Frickenhausen, Germany) and incubated with 150 µL *E. coli* TG1 (OD_600_ = 0.5) firstly for 30 min at 37°C, then 30 min at 37°C and 650 rpm to infect the phage particles. 1 mL 2xYT-GA (1.6% (w/v) Tryptone; 1% (w/v) Yeast extract; 0.5% (w/v) NaCl (pH 7.0), 100 mM D-Glucose, 100 µg/mL ampicillin) was added and incubated for 1 h at 37°C and 650 rpm, followed by addition of 1×10^10^ cfu M13KO7 helper phage. Subsequently, the infected bacteria were incubated 30 min at 37°C followed by 30 min at 37°C and 650 rpm before centrifugation for 10 min at 3220xg. The supernatant was discarded and the pellet resuspended in fresh 2xYT-AK (1.6% (w/v) Tryptone; 1% (w/v) Yeast extract; 0.5% (w/v) NaCl (pH 7.0), 100 µg/mL ampicillin, 50 µg/mL kanamycin). The antibody phage were amplified overnight at 30°C and 650 rpm and used for the next panning round. In total four panning rounds were performed. In each round, the stringency of the washing procedure was increased (20x in panning round 2, 30x in panning round 3) and the amount of antigen was reduced (2.5 µg in panning round 2, 1.5 µg in panning round 3). After third panning round single clones were analyzed for production of RBD specific scFv by screening ELISA.

#### Screening of monoclonal recombinant binders using E. coli scFv supernatant

Soluble antibody fragments (scFv) were produced in 96-well polypropylene MTPs (U96 PP, Greiner Bio-One) as described before (Russo et al., 2018; Wenzel et al., 2020a). Briefly, 150 μL 2xYT-GA was inoculated with the bacteria bearing scFv expressing phagemids. MTPs were incubated overnight at 37°C and 800 rpm in a MTP shaker (Thermoshaker PST-60HL-4, Lab4You, Berlin, Germany). A volume of 180 μL 2xYT-GA in a PP-MTP well was inoculated with 20 μL of the overnight culture and grown at 37°C and 800 rpm for 90 minutes (approx. OD_600_ of 0.5). Bacteria were harvested by centrifugation for 10 min at 3220xg and the supernatant was discarded. To induce expression of the antibody genes, the pellets were resuspended in 200 μL 2xYT supplemented with 100 μg/mL ampicillin and 50 μM isopropyl-beta-D-thiogalacto-pyranoside (IPTG) and incubated at 30°C and 800 rpm overnight. Bacteria were pelleted by centrifugation for 20 min at 3220xg and 4°C.

For the ELISA, 100 ng of antigen was coated on 96 well microtiter plates (High Binding, Costar) in PBS (pH 7.4) overnight at 4°C. After coating, the wells were blocked with 2% MPBST for 1 h at RT, followed by three washing steps with H_2_O and 0.05% Tween20. Supernatants containing secreted monoclonal scFv were mixed with 2% MPBST (1:2) and incubated onto the antigen coated plates for 1 h at 37°C followed by three H_2_O and 0.05% Tween20 washing cycles. Bound scFv were detected using murine mAb 9E10 which recognizes the C-terminal c-myc tag (1:50 diluted in 2% MPBST) and a goat anti-mouse serum conjugated with horseradish peroxidase (HRP) (A0168, Sigma) (1:42000 dilution in 2% MPBST). Bound antibodies were visualized with tetramethylbenzidine (TMB) substrate (20 parts TMB solution A (30 mM Potassium citrate; 1% (w/v) Citric acid (pH 4.1)) and 1 part TMB solution B (10 mM TMB; 10% (v/v) Acetone; 90% (v/v) Ethanol; 80 mM H_2_O_2_ (30%)) were mixed). After stopping the reaction by addition of 1 N H_2_SO_4_, absorbance at 450 nm with a 620 nm reference was measured in an ELISA plate reader (Epoch, BioTek). Monoclonal binders were sequenced and analyzed using VBASE2 (www.vbase2.org) (Mollova et al., 2010) and possible glycosylation positions in the CDRS were analyzed according to Lu et al (Lu et al., 2019).

#### ScFv-Fc and IgG production

Unique scFv sequences isolated by antibody-phage display were subcloned into pCSE2.7-hIgG1-Fc-XP using NcoI/NotI (New England Biolabs, Frankfurt, Germany) for mammalian production in Expi293F cells as scFv-Fc (Wenzel et al., 2020a). For IgG production, the variable domains were recloned into the IgG vectors human IgG1 format by subcloning of VH in the vector pCSEH1c (heavy chain) and VL in the vector pCSL3l/pCSL3k (light chain lambda/kappa) (Steinwand et al., 2014) adapted for Golden Gate Assembly procedure with Esp3I restriction enzyme (New England Biolabs, Frankfurt, Germany). For the antibodies CB6 (pdb 7C01), CR3022 (pdb 6W41) and REGN10933 and REGN10987 (pdb 6XDG) the public available amino acid sequences were used and the V-Genes were ordered as GeneArt Strings DNA fragments (Thermo Fisher Scientific, Schwerte, Germany) and recloned in the above indicated vectors. A “silenced” Fc part with following point mutations described Armour et al (Armour et al., 1999) and Shields et al (Shields et al., 2001) were used: E233P, L234V, L235A, deletion of G236, D265G, A327Q and A330S. Expi293F cells were cultured at 37°C, 110 rpm and 5% CO_2_ in Gibco FreeStyle F17 expression media (Thermo Fisher Scientific) supplemented with 8 mM Glutamine and 0.1% Pluronic F68 (PAN Biotech). At the day of transfection cell density was between 1.5 - 2×10^6^ cells/mL and viability at least above 90%. For formation of DNA:PEI complexes 1 µg DNA/mL transfection volume and 5 µg of 40 kDa PEI (Polysciences) were first diluted separately in 5% transfection volume in supplemented F17 media. DNA (1:1 ratio of the vectors for IgG production) and PEI was then mixed and incubated ∼25 min at RT before addition to the cells. 48 h later the culture volume was doubled by feeding HyClone SFM4Transfx-293 media (GE Healthcare) supplemented with 8 mM Glutamine. Additionally, HyClone Boost 6 supplement (GE Healthcare) was added with 10% of the end volume. One week after transfection supernatant was harvested by 15 min centrifugation at 1500xg.

#### Inhibition of S1-S2 and RBD binding to ACE2 expressing cells using flow cytometry

The inhibition tests in flow cytometry on EXPI293F cells were performed based on a previously published protocol (Bertoglio et al., 2020). Briefly, Expi293F cells were transfected according to the protocol above using pCSE2.5-ACE2_fl_-His and 5% eGFP plasmid. Two days after transfection, purified S1-S2-His was labelled using Monolith NT_TM_ His-Tag Labeling Kit RED-tris-NTA (Nanotemper) according to the manufacturer’s protocol. In this setup 50 nM antigen was incubated with min. 1 µM of different scFv-Fc and the ACE2 expressing cells. The resulting median antigen fluorescence of GFP positive living single cells was measured. For comparison of the different scFv-Fc first the median fluorescence background of cells without antigen was subtracted, second it was normalized to the antigen signal where no antibody was applied. ScFv-Fc showing an inhibition in this first setup were further titrated as IgGs (max. 500 nM-0.5 nM) on S1-S2-His, S1-His or on RBD-mFc (max. 100 nM-0.1 nM). S1-His and the corresponding mutants were detected with mouse anti-penta His (Qiagen) and goat anti-mFc APC-conjugated antibody (Dianova). RBD-mFc was detected directed with the goat anti-mFc APC-conjugated antibody (Dianova). Measurements were performed with MACSQuant Analyzer (Milteny Biotech). The IC_50_ was calculated using the equation f(x)=Amin+(Amax-Amin)/(1+(x0/x)^h)^s and parameters from OriginPro (2019).

#### Dose dependent binding of IgG in titration ELISA

For titration ELISA, purified IgGs were titrated from 3.18 μg/mL- 0.001 ng/mL on 30 ng/well of the following antigens: S1-S2-His (High Five cell produced), RBD-mFc (High Five cell produced), S1-mFc (High Five cell produced) and TUN219-2C1-mFc (as control for unspecific Fc binding). In addition, all scFv-hFc were also tested only at the highest concentration (3.18 μg/mL) for unspecific cross-reactivity on Expi293F cell lysate (10^4^ cells/well), BSA (1% w/v) and lysozyme. IgGs were detected using goat-anti-hIgG(Fc)- HRP (1:70000, A0170, Sigma). Titration assays were performed using 384 well or 96 well microtiter plates (Greiner Bio-One) using Precision XS microplate sample processor (BioTek), EL406 washer dispenser (BioTek) and BioStack Microplate stacker (BioTek). EC50 were calculated with by GraphPad Prism Version 6.1, fitting to a four-parameter logistic curve. Titration ELISAs on other coronaviruses and S1-HIS mutants were performed as described above.

#### Screening and titrating monoclonal antibodies for SARS-CoV-2 neutralization in cell culture

VeroE6 cells (ATCC CRL-1586) were seeded at a density of 6*10^4^/well onto cell culture 96-well plates (Nunc, Cat.#167008). Two days later, cells reached 100% confluence.

For titration, antibodies were diluted in 1/√10 steps and mixed with a fixed inoculum of SARS-CoV-2/Münster/FI110320/1/2020 (kind gift of Stephan Ludwig, University of Münster, Germany) (10-20, respectively 100-150 pfu) in a total volume of 500 µl of Vero E6 medium (DMEM, 10% FCS, 2 mM glutamine, penicillin, streptomycin). After one hour incubation at 37°C, cells were infected with the antibody/virus mix, incubated for one hour and then overlaid with Vero E6 medium containing 1.5% methyl-cellulose. Three days postinfection, wells were imaged using a Sartorius IncuCyte S3 (4x objective, whole-well scan) and plaques were counted from these images. Image data was quantified with the IncuCyte S3 GUI tools measuring the decrease of confluence concomitant with the cytopathic effect of the virus in relation to uninfected controls and controls without antibody and analyzed with Origin using the Logistic5 fit.

#### Specificity assay

To test specificity of the antibody candidates an ELISA on DNA, LPS, lysozyme and cell lysate was performed under standard conditions (see above). In brief, 10 µg/mL of the respective antigen was immobilized on 96 well microtiter plates (High binding, Costar) in PBS (pH 7.4) overnight at 4°C. After blocking 10 µg/mL of STE90-C11 IgG, Avelumab, Palivizumab and IVIG respectively were incubated and later detected using goat-anti-hIgG(Fc)-HRP (1:70000, A0170, Sigma). TMB reaction took place for 30 min and absorbance at 450 nm with a 620 nm reference was measured in an ELISA plate reader (Epoch, BioTek). All signals were normalized to the absorbance of Avelumab.

#### Analytical size exclusion chromatography (SEC)

All purified antigens and indicated antibodies were run on Superdex 200 Increase 10/300GL column (Cytiva) on Äkta pure system (Cytiva) according to the manufactures protocol.

#### Affinity measurement by Bio-Layer Interferometry

The affinity was measured by Bio-Layer Interferometry in three different assays using the Octet qKe (Fortebio/Sartorius GmbH, Göttingen, Germany).

In the first assay, anti-Mouse Fc-Capture (AMC) sensors were activated for 10 min in PBS. After that, the sensors were equilibrated in assay buffer (PBS containing 1% BSA and 0.05% Tween 20) for 60 s before RBD-mFc (Sino Biologicals) was loaded onto the sensors at 10 µg/ml for 180 s. After a stable baseline measurement was established (60 s), antigen-loaded sensors were transferred to an 8-point antibody dilution series (500, 150, 50, 15, 5, 1.5, 0.5 and 0 nM). Association of the Fab antibody to the antigen was measured for 300 s. After that, the sensors were transferred into assay buffer were the dissociation was measured for 900 s. Significant binding of the antibody to an unloaded sensor was not detected. For data analysis, the reference measurement (0 nM) was subtracted from the other measurements and data traces ranging from 150 to 1.5 nM were used for modeling of the kinetic data using a 1:1 binding model.

In the second assay, anti-human Fab (FAB2G) sensors were activated for 10 min in PBS. After that, the sensors were equilibrated in assay buffer (PBS containing 1% BSA and 0.05% Tween 20) for 60 s before the IgG antibody was loaded onto the sensors at 2.5 µg/ml for 180 s. After a stable baseline measurement was established (60 s), antibody-loaded sensors were transferred to an 8-point S1-HIS antigen dilution series (500, 150, 50, 15, 5, 1.5, 0.5 and 0 nM). Association of the S1 antigen to the antibody was measured for 300 s. After that, the sensors were transferred into assay buffer were the dissociation was measured for 900 s. Significant binding of the antigen to an unloaded sensor was not detected. For data analysis, the reference measurement (0 nM) was subtracted from the other measurements and data traces ranging from 50 to 5 nM were used for modelling of the kinetic data using a 1:1 binding model in the Data Analysis HT 11.0 software tool.

In the third assay, protein A sensors were activated for 10 min in PBS. Before use, the sensors were regenerated for 5 cycles in 10 mM Glycine buffer (pH2.0) followed by neutralization in PBS. Each step was performed for 5 s. After that, the regenerated sensors were equilibrated in assay buffer (PBS containing 1% BSA and 0.05% Tween 20) for 60 s before the IgG antibody was loaded onto the sensors at 2.5 µg/ml for 180 s. After a stable baseline measurement was established (60 s), antibody-loaded sensors were transferred to an 8-point S1-HIS antigen dilution series (500, 150, 50, 15, 5, 1.5, 0.5 and 0 nM). Association of the S1 antigen to the antibody was measured for 300 s. After that, the sensors were transferred into assay buffer were the dissociation was measured for 900 s. Significant binding of the antigen to an unloaded sensor was not detected. For data analysis, the reference measurement (0 nM) was subtracted from the other measurements and data traces ranging from 50 to 0.5 nM were used for modeling of the kinetic data using a 1:1 binding model.

#### Immunoblot analysis

For the immunoblot analysis RBD-His was disrupted either by incubation for 10 minutes at 56°C without β-Mercaptoethanol, at 95°C without β-Mercaptoethanol or at 95°C in Laemmli sample buffer (Laemmli, 1970) with β-Mercaptoethanol. RBD was separated by 12% SDS-PAGE and blotted onto a nitrocellulose membrane (Amersham Protan 0.2 µm NC, GE HealthCare). The membrane was blocked with 2% M-PBST for 1 h at RT. For the detection of RBD-His, 10 µg/mL STE90-C11 IgG was used. For control, RBD-His (95°C + β-Mercaptoethanol) was detected with 2µg/mL mouse anti-His (Dia-900-200, Dianova, Hamburg, Germany) for 1.5 h at RT. After 3x washing with PBST the secondary antibody goat anti-human Fc AP conjugated (1:20,000, Jackson ImmunoResearch, Cambridge House, UK) was used for the detection of STE90-C11 and goat anti-mouse IgG AP conjugated (1:30,000, 115-055-071, Dianova) was used for the anti-His antibody and incubated for 1 h. Finally, the membrane was washed 2x with PBST and 4x with PBS. Staining and visualization of specific proteins was performed with 5-Brom-4-Chlor-3-Indolyl-Phosphat/Nitro Tetrazolium Blue Chloride (NBT-BCIP, Thermo Fisher Scientific GmbH, Dreieich, Germany) according to standard protocols.

#### Hamster model of SARS-CoV-2 infection

Animal procedures were performed according to the European Guidelines for Animal Studies after approval by the Institutional Animal Care Committee and the relevant state authority (Landesamt für Gesundheit und Soziales, Berlin, Permit number 0086/20). SARS-CoV-2 isolate BetaCoV/Germany/BavPat1/2020 (Wölfel et al., 2020) was used as challenge virus for hamster experiments. The virus was propagated and titrated on Vero E6 cells (ATCC CRL-1586) in minimal essential medium (MEM; PAN Biotech, Aidenbach, Germany) supplemented with 10% fetal bovine serum (PAN Biotech), 100 IU/ml penicillin G and 100 µg/ml streptomycin (Carl Roth, Karlsruhe, Germany) and stored at −80°C prior to experimental infections.

Per group, nine male and female Syrian hamsters (Mesocricetus auratus) strain RjHAN:AURA (Janvier, Le Genest-Saint-Isle, France) were used. Animals were housed in GR-900 IVC cages (Tecniplast, Buguggiate, Italy) and provided with food ad libidum and bountiful enrichment and nesting materials (Carfil, Oud-Turnhout, Belgium). Hamsters were randomly distributed into experimental groups and treated intraperitoneally with 3.7 mg/kg or 37 mg/kg STE90-C11 in a total volume of 1 ml PBS, two hours post infection, the control group received 1 ml PBS only at the same time-point.

SARS-CoV-2 infection was performed as previously described (Osterrieder et al., 2020). Briefly, anaesthetized hamsters received 1×105 pfu SARS-CoV-2 in 60 µL MEM intranasally two hours before treatment. Following infection, the clinical presentation of all animals was monitored twice a day, body weight of all hamsters was recorded daily. On days 3, 5 and 14 post infection, three randomly assigned hamsters per group were euthanized. Euthanasia was applied by exsanguination under general anaesthesia as described (Nakamura et al., 2017). Oropharyngeal swabs and lungs were collected for virus titrations, RT-qPCR and/or histopathological examinations. All organs were immediately frozen at -80 °C or preserved in 4% formaldehyde for subsequent in-depth histopathological investigations.

To assess virus titers from 50 mg lung tissue, tissue homogenates were prepared using a bead mill (Analytic Jena) and 10-fold serial dilutions were prepared in MEM, and plated on Vero E6 cells in 12-well-plates. The dilutions were removed after 2 h and cells were overlaid with 1.25% microcrystalline cellulose (Avicel) in MEM supplemented with 10% FBS and penicillin/streptomycin. Three days later, cells were formalin-fixed, stained with crystal violet, and plaques were counted.

#### Transgenic mice model of SARS-CoV-2 infection

All animal experiments were performed in compliance with the German Animal Welfare Act (TierSchG BGBl. I S. 1206, 1313; May 18, 2006) and Directive 2010/63/EU. The mice were handled in accordance with good animal practice as defined by the Federation for Laboratory Animal Science Associations and Gesellschaft für Versuchstierkunde/Society of Laboratory Animal Science. All animal experiments were approved by the responsible state office (Lower Saxony State Office of Consumer Protection and Food Safety) under permits number 20_3567. K18hACE2 mice (B6.Cg-Tg(K18-ACE2)2Prlmn/J) were purchased from Charles River (Sulzfeld, Germany), Mice were housed at the animal facility of the Helmholtz Centre for Infection Research under pathogen-free conditions.

Mice were fixed in a restrainer before injection and the lateral tail veins were hyperaemized. Different antibody concentrations were diluted in 100ul of PBS and injected intravenous into the lateral tail vein.

Female and male at least 6-wk-old mice were infected with tissue culture– derived virus and housed in specific pathogen-free conditions throughout the experiment. Mice were anesthetized with Ketamin/Xylazin and inoculated intranasally with 2,000 PFU of virus in 20 ul of Phosphate buffered saline (PBS).

The mice were sacrificed by CO_2_ asphyxiation on day 5. Lungs were collected aseptically, homogenized in 500 µL PBS and stored at −80◦C. Part of the organ homogenates were used for titration cells and the other part for qPCR Analysis. Organ homogenates were serially diluted 1:10 – 1:10^5^ in medium (DMEM supplemented with 5% FCS, 2 mM glutamine, 100 IU/mL penicillin and 100 µg/mL streptomycin). Vero E-6 cells were then inoculated with 200ul of diluted homogenates and incubated for 1h, 37°C, CO_2_ incubator. Cells were then coverd with 1.75% Carboxymethyl-cellulose and incubated at 37°C, CO_2_ incubator for 3-5 days. Plates were then fixed with 6% Paraformaldehyde for 1h and then stained with 1% Crystal violet. Plaques were then read under microscope.

#### In vitro evolution of SARS-CoV-2 by antibody co-cultivation

This assay was performed with STE90-C11 and Palivizumab as described by Baum et al. (Baum et al., 2020).

#### Fab and RBD22 production for co-crystallization

The production of STE90-C11 Fab fragment was done by transient co-transfection of plasmids encoding the heavy and the light chain in Expi293F cells cultivated at 37°C, 5% CO2 and 100rpm in Expi Expression Medium. The transfection was performed at a density of 3*10^6 cells/ml by adding 1µg/ml culture mixed plasmids and 4 µg/ml culture PEI 40 kDa (Polyscience). The culture was incubated for 72 hours according to the protocol. The supernatant was harvested by centrifugation (30min, 3000g) and sterile filtration (0.2µm). The Fab-fragment was purified by affinity chromatography using 1ml HisTrap Excel column (GE Healthcare) followed by a size exclusion chromatography on a 10/300 Superdex200 Increase column (GE Healthcare) according to manufactures manual.

For the production of RBD22 High Five cells grown in EX-CELL 405 serum-free medium (Sigma) were transiently transfected with 5µg/mL plasmid and 20µg/mL PEI 40 kDa (Polyscience) at a cell density of 5*10^6 cells/ml. After 4h incubation at 27°C and 100rpm the cells were diluted 5-fold with EX-CELL medium to a density of 1*10^6 cells/ml and further incubated for three more days. Following centrifugation (5000xg 30min) and sterile filtration (0.2µm) the supernatant was loaded on a 5ml HisTrap Excel column (GE Healthcare). After washing and elution with imidazole the RBD22 containing fractions were further purified by size exclusion chromatography on a 26/600 Superdex 200 pg using 20mM Tris-HCl pH8 and 150mM NaCl as a buffer. Aliquots were snap-frozen in liquid nitrogen and stored at -80°C till further use.

#### Crystallization, data collection and structure determination

The Fab fragment of STE90-C11 was incubated overnight with a 1.2 molar excess of purified RBD22 at 4°C. The complex was isolated by size exclusion chromatography on a Superdex 200 Increase 10/300GL (GE Healthcare) with 20mM Tris-HCl pH8 and 150mM NaCl as a running buffer. Fractions containing the complex were concentrated utilizing a Vivaspin2 ultrafiltration unit (10,000 MWCO; Sartorius). Crystallizations trails were set up in 96-well sitting-drop vapor diffusion plates (Intelli 96-3 plates, Art Robbins Instruments) with a pipetting robot (Crystal Gryphon, Art Robbins Instruments) mixing 200nl reservoir solution with 200nl of protein solution (14mg/mL, 7mg/mL) and equilibrated against 60 μl of reservoir solution. As initial screens the Cryos Suite and the JCSGplus Suite (Qiagen) were chosen and crystal growth was monitored in a crystal hotel (RockImager, Formulatrix). Initial hits were further optimized by a random screen assembled with a Formulator pipetting robot (Formulatrix). Best diffracting crystals grew in 13.3% (w/v) polyethylene glycol 6,000, 0.1M MES pH 5.6 and 0.24M tri sodium citrate. Crystals were harvested with nylon loops and soaked in reservoir solution mixed with 2,3-(R,R)-butandiol (Alfa Aeser) to a final concentration of 10% (v/v) prior to flash cooling in liquid nitrogen.

3600 diffraction images with an oscillation angle of 0.1° per image were collected at the beamline P11 at PETRA III (DESY, Hamburg, Germany) (Burkhardt et al., 2016) on a Pilatus 6M fast detector (Dectris) and processed with XDS (Kabsch, 2010) and Aimless (Evans and Murshudov, 2013) yielding a dataset with a resolution cut off of 2.0 Å based on a CC1/2 value greater than 0.5. Initial phases were determined by molecular replacement with Phaser (McCoy et al., 2007). As a search model the coordinates of a Fab fragment and the RBD was used (PDB: 7BWJ) (Ju et al., 2020). The model was further improved by manual rebuilding in Coot (Emsley et al., 2010) and computational refinement with phenix. refine (Afonine et al., 2012) including placement of water, TLS refinement and riding hydrogens in the final steps of the procedure. Depictions of the model were generated with PyMol molecular graphics system (Schrödinger LLC; version 2.3.2). Data processing and model refinement statistics can be found in Supplementary Data 11. The final model can be accessed under the PDB code 7B3O.

## Acknowledgements

We kindly acknowledge the financial support of MWK Niedersachsen (14-76103-184 CORONA-2/20) for the projects “Antibody generation”, “Neutralization experiments” and “Structure-based analysis of antiviral strategies against CoV-2 target proteins”. We also acknowledge support of the European Union for the ATAC (“antibody therapy against corona”, Horizon2020 number 101003650) consortium. We acknowledge DESY (Hamburg, Germany), a member of the Helmholtz Association HGF, for the provision of experimental facilities. Parts of this research were carried out at PETRAIII and we would like to thank Johanna Hakanpää for assistance in using the P11 beamline. We are deeply grateful to Adelheid Langner, Andrea Walzog, Bettina Sandner, Cornelia Oltmann and Wolfgang Grassl for constant help and support during the pandemic.

## Author contributions

F.B., A.F., T.S., S.D., J.v.d.H. L.Ĉ-Ŝ., M.S., M.H. conceptualized the study. F.B., V.F., M.R., P.A.H, L.A., U.R., T.K., D.M., N.L, S.S., R.B., K-T.S., K.D.R.R., P.R., D.S., A.K., S.Z.-E., Z.H., K.E., Y.K., M.B., Marg.S., G.M.S.G.M., E.V.W., G.R., J.A., J.T., M.S. performed and designed experiments. F.B., D.M. N.L., S.S., U.R., L.S., P.K., J.v.d.H., J.T., A.F., L.Ĉ-Ŝ., M.S., M.H. analyzed data. H.S.P.G., S.C., A.G. provided patient samples and support. S.D., L.Ĉ-Ŝ., M.H. conceived the funding. P.K., E.V.W., Gi.R., A.H., V.F., S.D., Gü.R., A.F., J.v.d.H., M.S., M.H. advised on experimental design and data analysis. F.B., T.K., S.D., L.Ĉ-Ŝ., M.S., M.H. wrote the manuscript.

## Declaration of interests

The authors declare a conflict of interest. F.B., D.M., N.L., S.S., P.A.H., R.B., M.R., K.T.S., K.D.R.R., S.Z.-E., M.B., V.F., S.T., M.S. and M.H. are inventors on a patent application on blocking antibodies against SARS-CoV-2. A.H., A.F. and T.S. are officers of CORAT Therapeutics GmbH, a company founded by YUMAB GmbH for the clinical and regulatory development of STE90-C11 (development name COR-101) (ClinicalTrials.gov ID: NCT04674566 for safety and efficacy of COR-101 in hospitalized patients with moderate to severe COVID-19). A.H. is shareholder of CORAT Therapeutics GmbH. S.D. and M.H. are advisors of Corat Therapeutics GmbH. A.F., T.S., S.D. and M.H. are shareholders of YUMAB GmbH.

## Notes

### Summary of Updates

The highlights of the revised version are: - binding and inhibition analysis of more RBD mutants including the binding and inhibition data for B.1.617 - data from two in vivo challenge models (hamster and mice)

